# Subcellular structure segmentation from cryo-electron tomograms via machine learning

**DOI:** 10.1101/2020.04.09.034025

**Authors:** Li Zhou, Chao Yang, Weiguo Gao, Talita Perciano, Karen M. Davies, Nicholas K. Sauter

## Abstract

We describe how to use several machine learning techniques organized in a learning pipeline to segment and identify subcellular structures from cryo electron tomograms. These tomograms are difficult to analyze with traditional segmentation tools. The learning pipeline in our approach starts from supervised learning via a special convolutional neural network trained with simulated data. It continues with semi-supervised reinforcement learning and/or a region merging techniques that try to piece together disconnected components that should belong to the same subcellular structure. A parametric or non-parametric fitting procedure is then used to enhance the segmentation results and quantify uncertainties in the fitting. Domain knowledge is used in generating the training data for the neural network and in guiding the fitting procedure through the use of appropriately chosen priors and constraints. We demonstrate that the approach proposed here work well for extracting membrane surfaces of protein reconstituted liposomes in a cellular environment that contains other artifacts.

## 1 Introduction

Despite the tremendous progress made in biological imaging that has yielded tomograms with ever-higher resolutions, the interpretation of data, (e.g., the segmentation of cell tomograms into organelles and proteins) remains a challenging task. The difficulty is most extreme, in our experience, in the case of cryo-electron tomography (cryo-ET), where the samples exhibit inherently low contrast due to the limited electron dose that can be applied during imaging before radiation damage occurs. The resulting tomograms thus have a low signal-to-noise ratio (SNR), as well as missing-wedge artifacts caused by the limited sample tilt range that is accessible during imaging [1]. Figure 1 shows one slice of a partial cryo-EM tomogram of membrane-bound proteins reconstituted into liposomes.

**Fig. 1.**
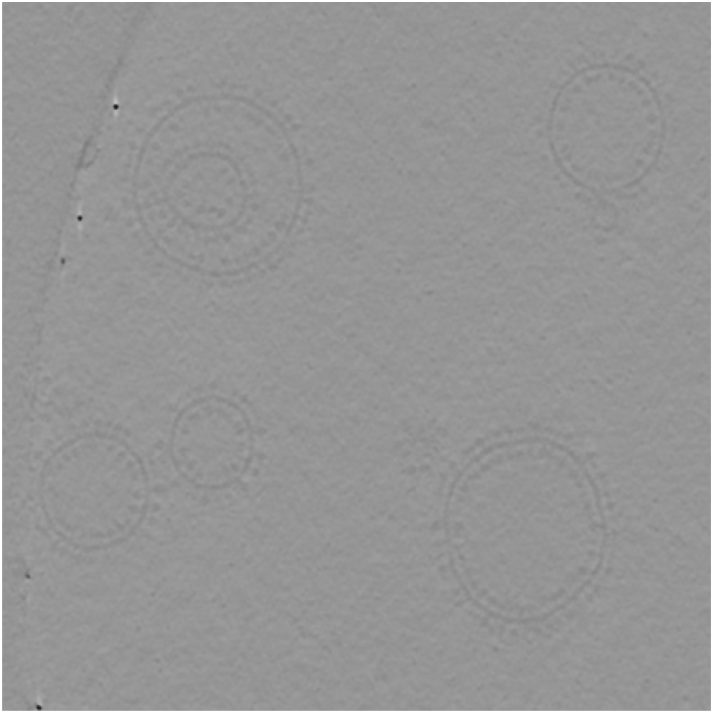
A slice of a partial cryo-EM tomograph of membrane-bound ATP synthase proteins reconstituted into liposomes. The proteins are shown as small dots attached to the circular-shaped liposome membranes.

Our objective is to identify and isolate from such tomograms multiple cellular substructures such as membranes, organelles and protein complexes that can be further analyzed. This objective is often achieved through an image segmentation procedure. Currently, such a procedure is performed, in most cases, by a human expert manually tracing or highlighting specific features in a tomogram, which are then extracted and analyzed for length, curvature, volume, distance, etc. This is an extremely time-consuming and labor-intensive process.

Although a number of automated segmentation algorithms and tools have been developed in the last few decades for high contrast medical 3D imaging [2-8], most of them perform poorly on cryo-ET datasets.

While SNR can be partially improved by applying contrast enhancement and edge detection algorithms, such as nonlinear anisotropic diffusion, wavelet transforms, or Sobel filters, these algorithms can also generate false connectivity and additional artifacts that degrade the results produced by automatic segmentation methods.

The reason why a human scientist can do a much better job at segmenting and extracting subcellular structures than a computer program is that he/she has prior knowledge (size, shape, etc.) about the biological object to be segmented. If we can train a machine to learn such knowledge, it may be possible to develop a more reliable automated segmentation tool that can be used to improve the throughput of the visualization and analysis and tie the structure to function, etc.

In recent years, there has been tremendous progress in the development of machine learning tools for image analysis and segmentation. In particular, convolutional neural network (CNN) based tools such as U-Net [9] have been developed for cryo-ET segmentation, where any arbitrary feature may be selected from the tomogram to be used as CNN training data [10]. Although the output is promising, this automatic machine learning algorithm still suffers from problems similar to pixel-based density thresholding algorithms used to assist manual segmentation. In addition, the success of this approach is hampered by the limited number of existing segmented structures to be used for training. Even though the recent development of cryo-electron tomography has produced many tomograms, high-quality substructure segmentations that can be used to train a neural network are still scarce and will always be scarce.

Given the complexity of the segmentation task and the inherent challenge in obtaining high-quality tomograms, it is unlikely a single image processing or machine learning technique can produce satisfactory results.

However, multiple machine learning techniques can be combined to enhance the segmentation results produced by a CNN based procedure. Among these are 1) reinforcement learning algorithms that can be used to connect multiple segmented pieces that belong to the same subcellular structure 2) classification algorithms that can separate different subcellular structures and place fragments of the same structure into the same group. 3) parametric and non-parametric fitting algorithms that produce a smooth and continuous surface representation of membranes.

In this paper, we will illustrate how these methods can be combined in an image analysis and segmentation pipeline that can significantly enhance the fidelity of segmentation of cryo-tomograms.

Although some of these methods can be directly applied to 3D tomograms, the large data volume of cellular cryo-tomograms makes direct 3D segmentation computationally costly in practice. Therefore, we choose to perform the 2D segmentation of tomogram slices first and refine these segmentation results in 3D by taking into account the correlation among images in adjacent slices of the tomogram.

This paper is organized as follows. In the next section, we will provide an overview of the main workflow of the overall segmentation procedure and how they fit together to meet the ultimate of structure analysis goal. This is followed by detailed discussion of each individual component of the methodology which includes the preprocessing of tomogram slices to improve image contrast (section 3), the initial segmentation by U-Net (section 4), the refinement of the segmentation in 2D using reinforcement learning, classification, and parametric/non-parametric fitting (section 5), as well as 3D refinement (section 6).

## 2 Main Workflow

Figure 2 depicts the overall workflow of the machine learning-based tomogram segmentation strategy we propose to analyze cryo-ET images. We first preprocess the tomogram slices to enhance the image contrast using the techniques to be presented in the next section.

**Fig. 2.**
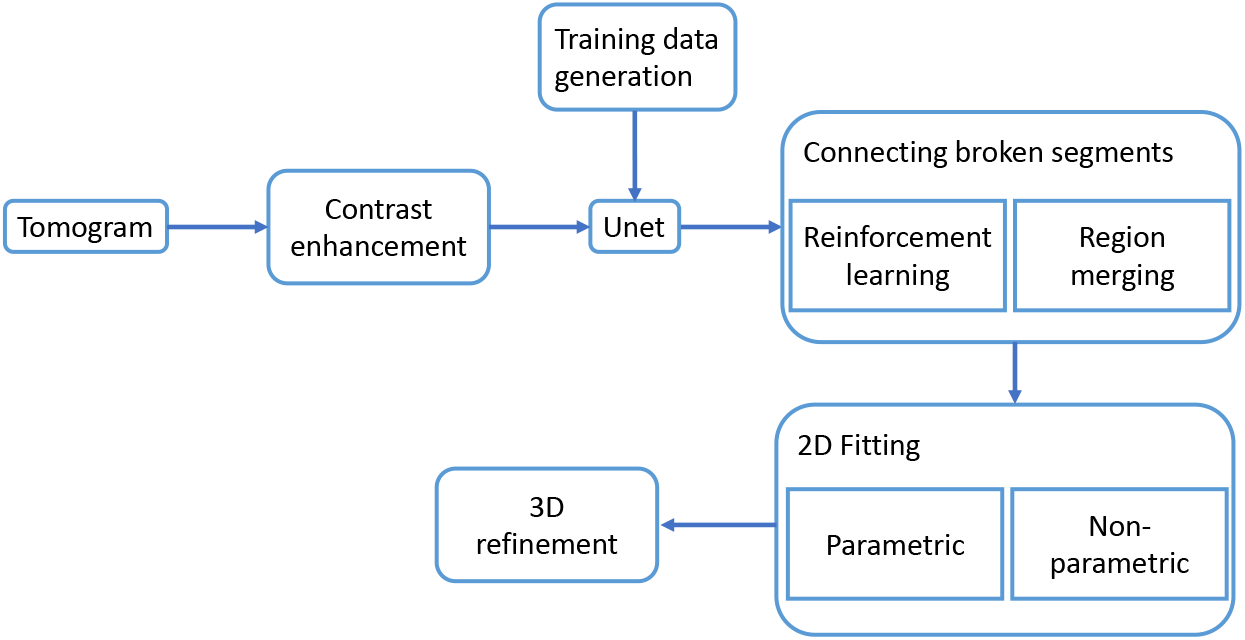
The main workflow of a machine learning-based approach that combines a number of techniques for segmenting cryo-EM tomograms and improving the segmentation.

We then generate training data for a U-Net by taking into account prior knowledge of the type of subcellular structure we plan to segment and analyze. The generation of the training data combines simple 2D geometric motifs with measured signal and noise features in the tomogram.

The training data is then used to train a U-Net, a CNN based segmentation tool that identifies subcellular structures that match the geometric motifs used in the training data from tomogram slices.

Because the output from the U-Net is typically not perfect and may contain fragmented components and artifacts, it is corrected by a 2D refinement procedure that tries to identify components that belong to the same subcellular structure using either a reinforcement learning algorithm or a region merging based algorithm.

A parametric or non-parametric nonlinear fitting procedure is then used to create smooth and continuous boundaries of membrane structures.

Corrected 2D sections are then combined in 3D and refined through a non-parametric fitting procedure to produce the final 3D segmentation.

## 3 Preprocessing by Contrast Enhancement

The preprocessing step used in our pipeline includes the application of the bilateral filter and an adaptive local contrast enhancement. The bilateral filter is an edge-preserving and noise-reducing filter [11]. It averages pixels based on their spatial closeness and radiometric similarity. In other words, it smooths homogeneous regions of the image and preserves details (such as borders of objects). After improving the signal-to-noise ration using the bilateral filter, the next step is to emphasize targeted structures for segmentation. We apply a technique called Contrast Limited Adaptive Histogram Equalization (CLAHE) [12]. This method uses histograms computed over different tile regions of the image. In doing so, local details are enhanced even in regions that are darker or lighter than most of the image. The final result after applying the two steps above to the image shown in Fig. 1 is presented in Fig. 3. In this image, the organelles in the tomogram, as well as the membrane-embedded ATP synthase proteins, are much more distinctive compared to the raw data, making this image more suitable for the segmentation step.

**Fig. 3.**
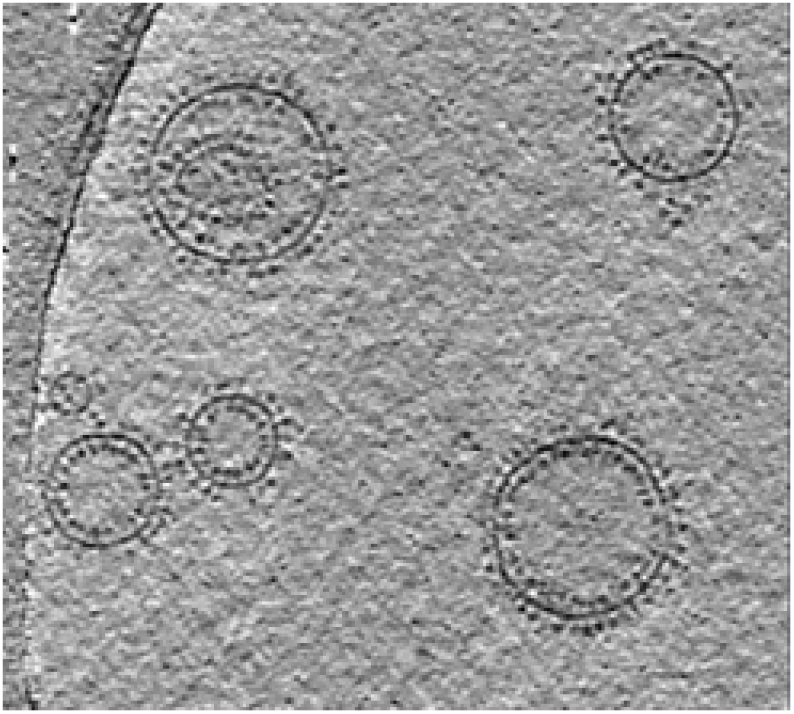
Contrast-enhanced tomogram slice

## 4 Segmentation by U-Net

U-Net [9] is a convolutional neural network (CNN) [13] based segmentation tool that has enjoyed tremendous success in biomedical image segmentation. The letter U in the name characterizes the layout of the CNN, which consists of a contracting path (the left half the U) and an expansion path (the right half of the U.) The contracting path maps the input image to a set of features through successive layers of convolution, rectified linear unit (ReLU) and max pooling operations. The expansive path upsamples the feature channels before convolving them with weighting matrices, concatenating with them feature maps produced in the contracting path, and feeding them into the ReLU layer.

One of the algorithmic ingredients that make U-Net robust is its ability to use excessive data augmentation generated by applying elastic deformations to the available training images. This algorithmic feature allows us to use a few well defined geometric motifs (such as circles and ellipses) to generate training data without relying on manual segmentated data that are difficult to obtain.

### 4.1 Generating training Data

Ideally, a U-Net should be trained by a set of manually segmented tomogram slices. However, because it is often time-consuming to perform manual segmentation, there is generally a limited amount of labeled data we can use. Hence it is not realistic to rely on using manually segmented tomograms to train a U-Net to perform additional segmentation.

Fortunately, training a U-Net does not necessarily require using precisely segmented images. We train the network to recognize subcellular structures that often have characteristic geometric features and shapes. If there is prior knowledge about the general shapes and features of these subcellular structures, we can generate training data through simulation.

The simulated 2D images we generate combine simple geometric motifs (such as ellipses and circles) with a simulated noisy background. The intensity profiles of both the geometric motifs and the background are chosen to match those in the tomogram to be segmented.

For example, Figure 4 shows the intensity of a selected region of the background and the histogram of the pixel intensities within this region, which clearly exhibits a Gaussian distribution. Figure 5 shows part of a liposome membrane with the background (a) and the mask used to extract membrane pixels (b). The histogram of membrane pixel intensity is also shown (c). Although the histogram does not strictly represent a Gaussian distribution, we can fit the histogram with a Gaussian. Gaussian fittings are performed on the histograms associated with both the background and the membranes to produce parameters we can use to generate simulated images.

**Fig. 4.**
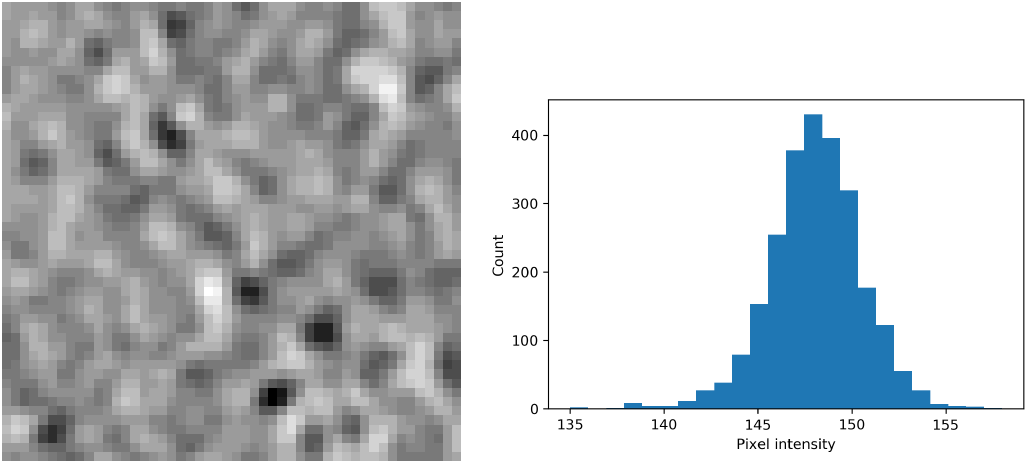
The histograms of the background intensity.

**Fig. 5.**
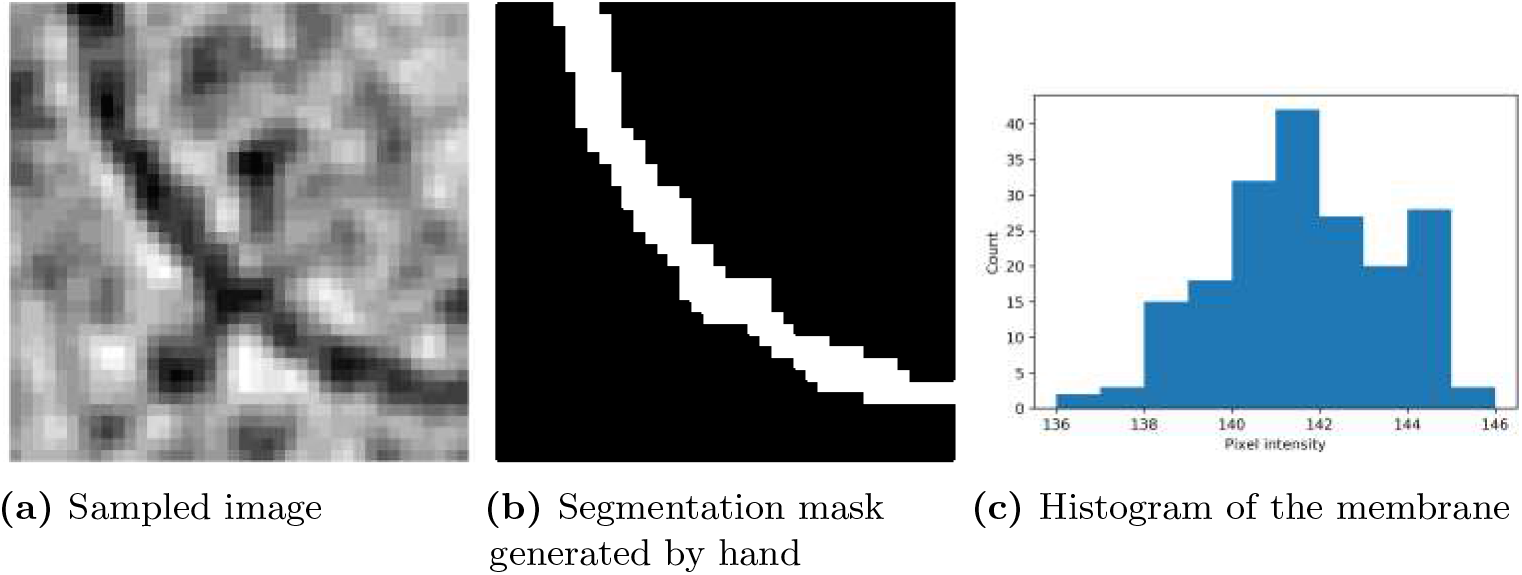
The histograms of the membrane intensity.

The simulated membranes we generated have different sizes and thicknesses. Their positions and orientations are randomly chosen. Figure 6 shows one of such simulated images.

In addition to the membranes, we also generate small solid circles near the membrane to mimic intrinsic membrane proteins (e.g., ATP synthase) with globular domains adjacent to the membrane. We label the membranes and proteins separately. The use of three distinct labels, i.e., 0 for background, 1 for the membrane and 2 for protein, (see Figure 6) significantly improves the segmentation result.

**Fig. 6.**
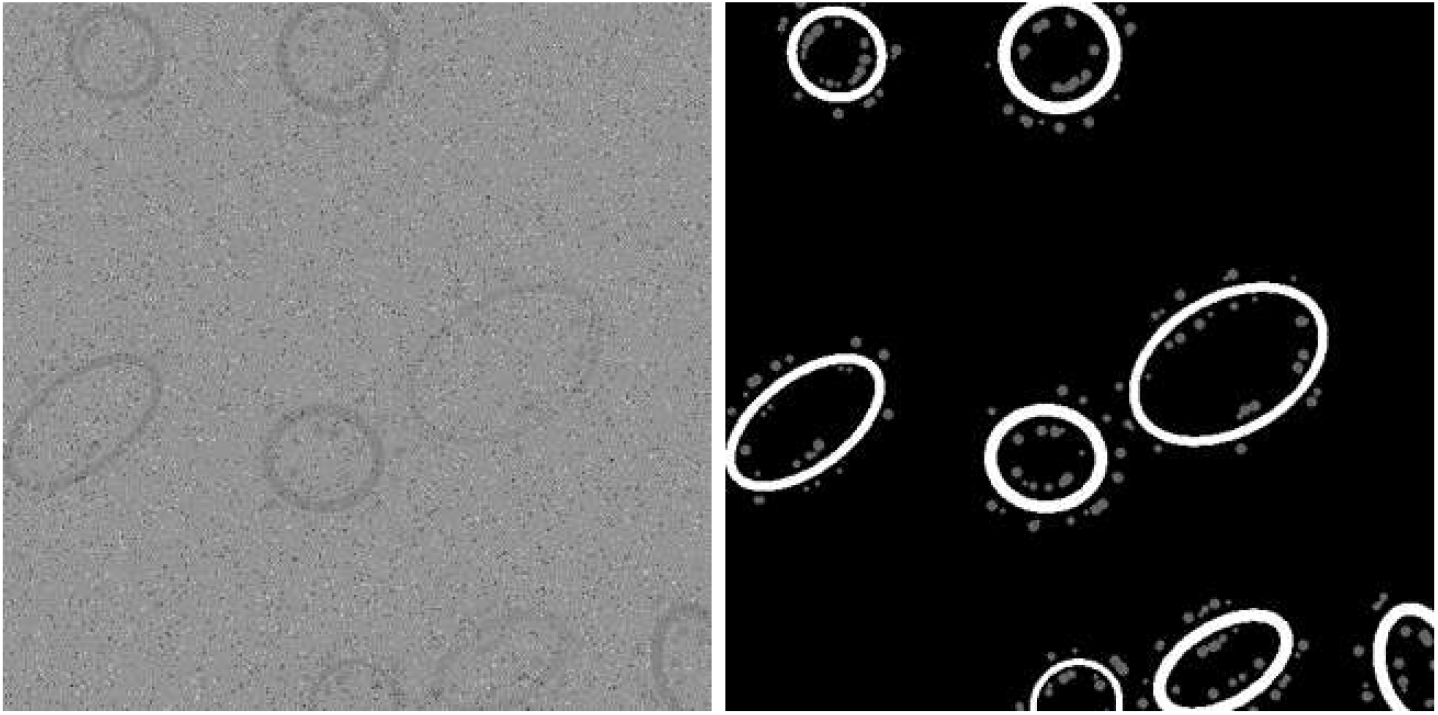
The simulated membranes and protein particles (left) and their labels (right).

### 4.2 Training and testing

To segment images like the one shown in Figure 1, we use 2000 simulated images to train the U-Net. For each input pixel *x*, the U-Net computes the activation function *a_k_*(*x*), which gives the likelihood of *x* being in the class *k*, for *k* = 0, 1, 2. A softmax function *p_k_*(*x*), which is defined as

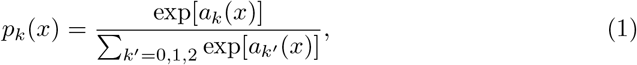

normalizes the likelihood to a value between 0 and 1. If *p_k_*(*x*) = 1, U-Net predicts *x* to be in class *k* with full certainty. If *p_k_*(*x*) = 0, U-Net predicts *x* not to be in class *k*.

Progress of the training process can be monitored by examining a loss function *E*, which is defined by the cross entropy

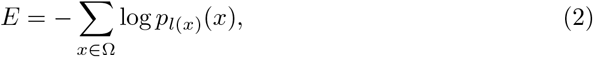

at each iteration (epoch), where *l*(*x*) is the true class label for pixel *x*, and Ω is the set of all pixels in the image to be segmented. A perfect segmentation would yield *E* = 0.

During the training process, an extra 5 images are used for testing, and 10 additional simulated images are reserved for validation after the training is over. Figure 7(a) shows the average loss function for 5 test images decreases rapidly as the number of training epochs increases. Figure 7(b) shows that the loss function associated with the validation images decreases in general also, but the change of *E* is not monotonic, and it is less smooth also.

**Fig. 7.**
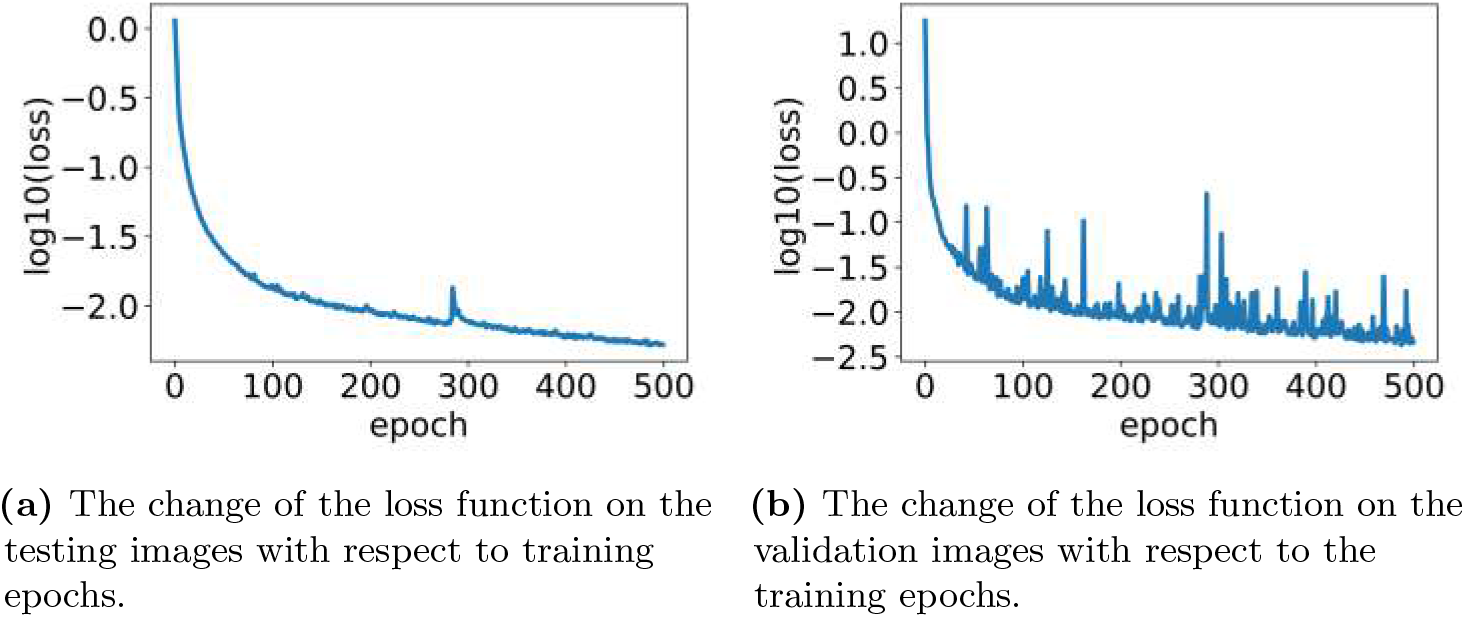
The convergence of the training process.

We choose the ADAM method [14] to train the U-Net by minimizing the loss function. The learning rate of the training method is set to 0.001, the exponential decay rate of the first and second moments is set to *β*_1_ =0.9 and *β*_2_ = 0.999 respectively. The training batch size is set to 20. The choice of these hyperparameters are usually problem-dependent, and may need to be optimized for different datasets.

Once the U-Net is trained, we can check the accuracy of the segmentation by computing the ratio of pixels assigned with the correct class labels and the total number of pixels in the image to be segmented. Table 1 shows the accuracy of the segmentation for 10 validation images in the first column. The overall classification accuracy is over 99%. This means more than 99% of pixels are correctly classified. In addition to this accuracy measurement, we also report the Intersection-Over-Union(IoU) value for each of the validation images. IoU is a widely used metric for evaluating the quality of image segmentation. It defined as

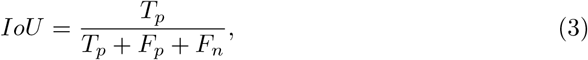

where *T_p_* is the number of true positives, *F_p_* is the number of false positives, and *F_n_* is the number of false negatives. IoU places more emphasis on the ratio between correctly classified feature pixels (True positives) over incorrectly classified pixels (false positives and false negatives). Column two in Table 1 gives the total IoUs for all pixels, which are less than the accuracy metric reported in Column 1. We can in fact evaluate IoU for each class (i.e., background, membrane and protein). These values are reported in columns 3–5. We can see that IoU values for background pixels are consistently higher than those of membrane and protein pixels. The membrane pixels seem to have the lowest IoU, indicating that it is more difficult to segment membranes from the background than the proteins.

**Table 1.** The accuracy of the U-Net segmentation on the validation images.

| Accuracy | Total IoU | IoU (background) | IoU (membrane) | IoU (protein) |
| --- | --- | --- | --- | --- |
| 0.996 | 0.949 | 0.995 | 0.885 | 0.968 |
| 0.996 | 0.949 | 0.995 | 0.885 | 0.968 |
| 0.996 | 0.942 | 0.996 | 0.865 | 0.966 |
| 0.995 | 0.946 | 0.994 | 0.882 | 0.962 |
| 0.997 | 0.950 | 0.997 | 0.893 | 0.961 |
| 0.995 | 0.949 | 0.995 | 0.883 | 0.968 |
| 0.995 | 0.948 | 0.995 | 0.874 | 0.974 |
| 0.997 | 0.954 | 0.997 | 0.899 | 0.966 |
| 0.996 | 0.953 | 0.996 | 0.894 | 0.969 |
| 0.994 | 0.941 | 0.994 | 0.877 | 0.952 |
| 0.995 | 0.945 | 0.995 | 0.871 | 0.968 |

### 4.3 U-Net segmentation results

Figure 8 shows the initial segmentation results produced by the U-Net for one of the tomogram slices shown in Figure 3. The liposome membranes shown in Figure 8(a) are separated from the proteins shown in Figure 8(b). These two types of subcellular structures are shown together in Figure 8(c).

**Fig. 8.**
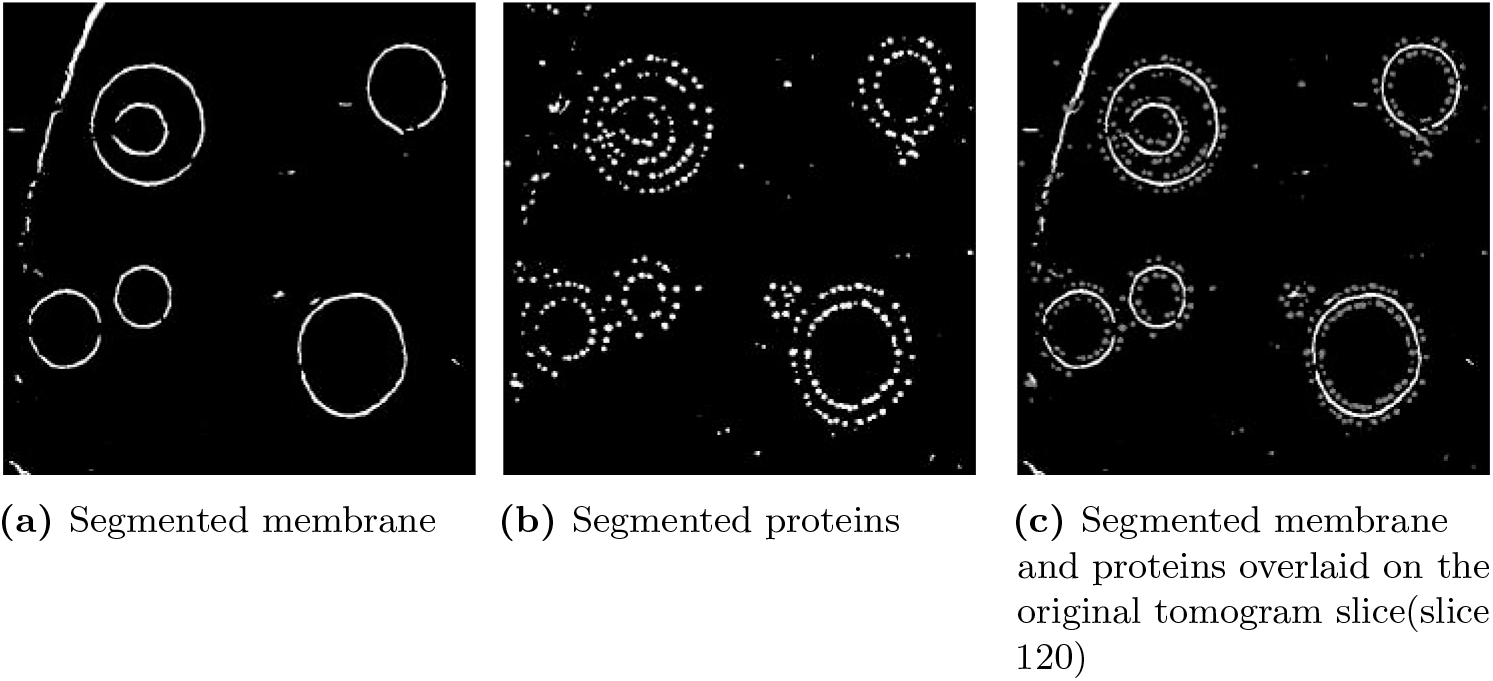
U-Net segmentation

## 5 Connecting Broken Segments

Although the U-Net does a remarkable job at identifying membranes of subcellular structures as shown in Figure 8, some of the membrane segments are disconnected. The gaps in the segmented membrane result from 1) low contrast and signal to noise ratio in the tomogram 2) incomplete tomogram reconstruction due to the missing wedge problem.

However, human vision can easily recognize how some of the disconnected components should be joined. With some prior knowledge of the possible shapes of the targeted subcellular structure, we can deduce how the disconnected components should be connected and how open boundaries can be closed. In this section, we discuss how to use a number of learning algorithms to join all disconnected membrane segments that should lie on the same subcellular membrane surface.

Our goal is to perform this type of postprocessing in an automated fashion with as little human intervention as possible. The challenge is that a tomogram may contain multiple subcellular structures, each enclosed by a membrane. If there were only one such structure, we could possibly use a curve or model fitting procedure to connect the membrane segments identified by the U-Net. Other contour completion algorithms may also be used [8].

In the presence of multiple subcellular structures, we need to determine, in an automated fashion, which labeled pixels belong to the same membrane segment, and which segments belong to the same membrane surface. While the first question is relatively easy to address by grouping labeled pixels that are within e-distance with each other, the second question is much harder to address because the labeled membrane segments can have different shapes, lengths, and curvatures, etc.

We present two strategies for achieving this task. The first strategy is based on reinforcement learning. The second strategy is based on region-based pixel merging.

### 5.1 Reinforcement Learning

Our first strategy is to train an agent to walk along segmented components and make connections with other segmented components with the goal of returning to the point it started from without crossing any segmented components that have already been traversed. Once the agent successfully returns to the starting point, the traversed segments are selected for further processing, and the agent can start again from a segmented component that has not been traversed. Otherwise, the agent is allowed to backtrack or start a new exploration trip (episode) if the existing journey is unlikely to be successful. The learning process is terminated when the number of attempts to traverse and return to the starting point exceeds a preset number. This type of learning algorithm is often referred to as *reinforcement learning*.

A reinforcement learning (RL) algorithm is often characterized by an agent performing a sequence of tasks to move from state to state in order to reach a certain goal. Each task involves taking one of the actions from a predefined set of actions. Each action is associated with a local reward used to indicate incremental progress toward accomplishing the final goal. Which action to take depends on a policy, which is described by what is often called a *Q*-value table (*Q*-table) that assigns a value to each (state, action) pair. For each state, the action that yields the largest *Q*-value is taken. The *Q*-table can be constructed iteratively by a training procedure in which the agent is encouraged to take an action that leads to the highest reward. However, randomness is built into the training algorithm so that the agent can take a locally non-optimal action from time to time at any given state to explore a larger search space. The stochastic nature of the training algorithm is formally characterized as a Markov decision process in which transition probabilities between different states and local reward for each transition are taken into account to define an iterative process that should ultimately yield the desired *Q*-table. However, in practice, due to the large number of states and the difficulty of defining transition probabilities in advance, the mapping between a (state, action) pair and its *Q*-value is constructed approximately by some means.

The RL algorithm we use consists of two phases. In the first phase, an agent is trained to traverse through U-Net segmented pixels that are sufficiently close with the goal of creating ordered lists of pixels that are connected. The walker starts at a labeled pixel on a segmented component that has not been included in any of the connected membrane surfaces. It moves around by taking one of the eight actions (move up, down, left, right, up then left, down left, up right, down right by one pixel).

The *Q*-table that defines the walking policy is constructed iteratively by taking into account the geometric features (position, orientation and curvature) of the segmented components as well as the local reward that is dynamically defined.

When the walker is on a segmented component, the reward for each of its actions is defined by the number of new labeled pixels resulting from the change in the field of view, which is defined to be a rectangular window of a certain size. Figure 9 shows that, when the walker moves from the blue pixel (state) to the red pixel (state), its new field of view (defined by the red rectangle) contains 4 new labeled pixels (enclosed by the yellow box in the image). This is the local award associated with the action that takes the walker from the blue pixel to the red pixel. In the case in which multiple actions yield the same reward, the action that keeps the walker in the middle of the segmented component is favored. This modified reward is also the Q-value associated with that particular (state, action) pair. By taking the actions that are associated with the largest *Q*-values, the walker traces out the segmented component as a 1D parameterized curve. When the walker reaches the end of a segmented component, it returns to the starting point and tries to traverse in the opposite direction if that has not been done.

**Fig. 9.**
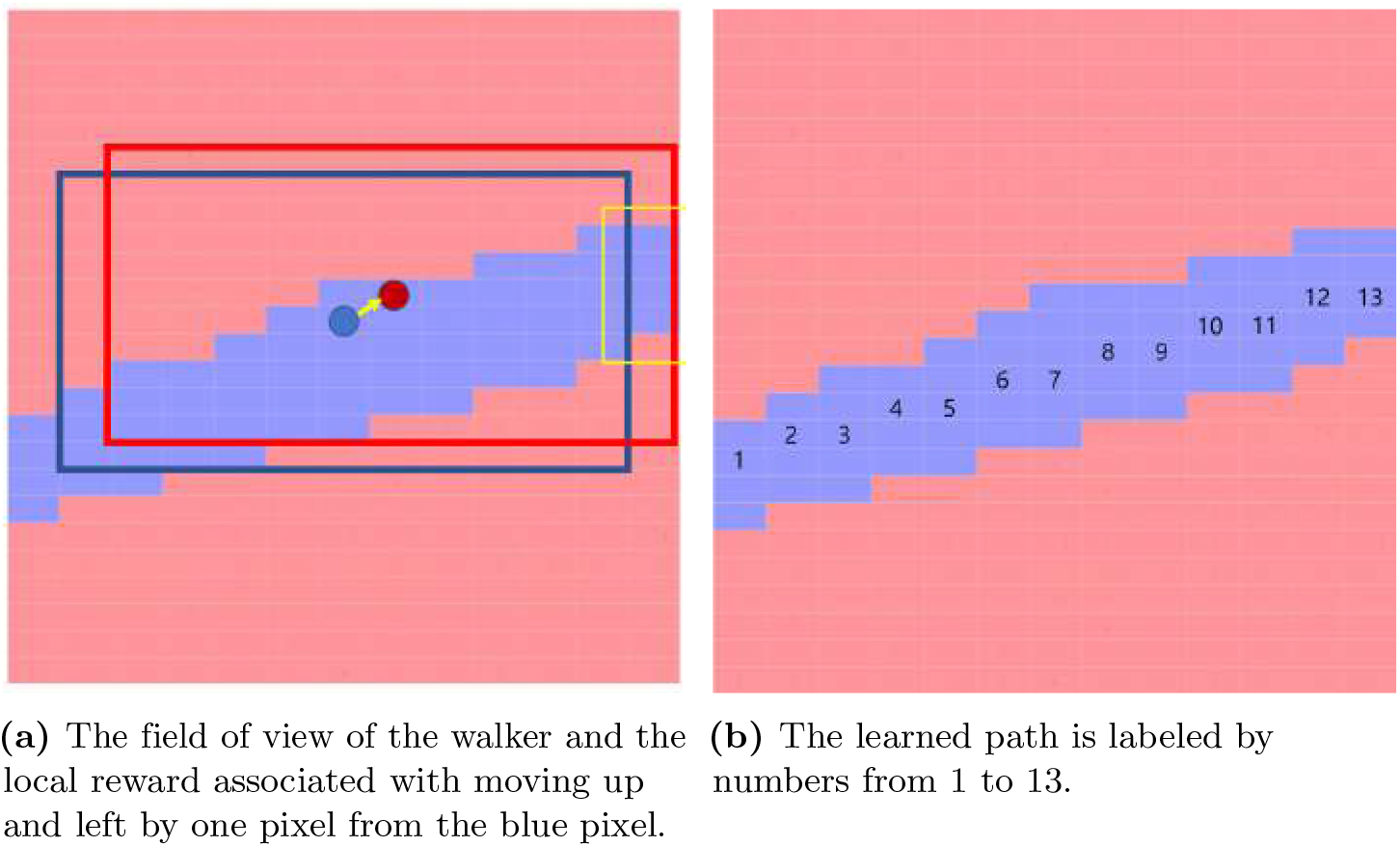
A schematic illustration of how local reward is calculated in the reinforcement learning algorithm as the walker traverses the segmented component (a) and the learned path (b).

Otherwise, it picks another pixel on another segmented component and repeats the same process until all segmented components have been traversed. This procedure allows us to generate a set of ordered and connected pixels (segments) from the output returned from the U-Net. We remove segments that have very few pixels. These tend to be introduced by the noise picked up by U-Net. Figure 10 shows all segments identified by the RL algorithm when it is applied to the U-Net output shown in Figure 8(a). Each segment is labeled by a number.

**Fig. 10.**
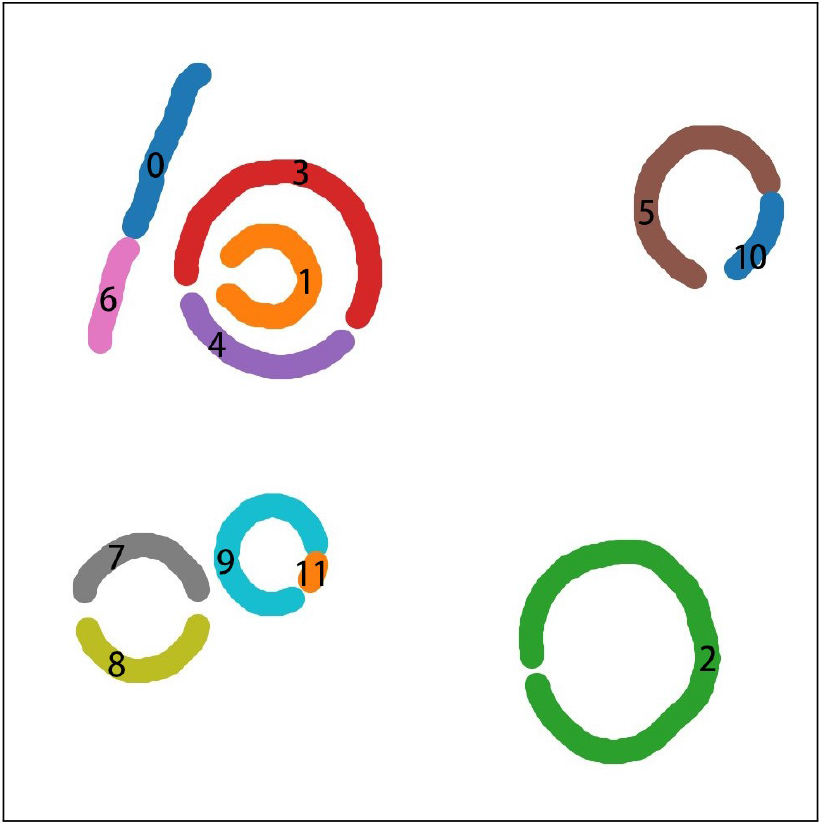
All connected segments identified by the RL algorithm after it is applied to the tomogram slice shown in Figure 8. Each segment is labeled by a distinct number.

In the second phase of the RL algorithm, our goal is to connect segmented components (ordered lists of pixels) identified in the first phase if they belong to the membrane of the same subcellular structure. The walker is initialized to exit from an endpoint *A* of a segmented component, e.g. as shown in Figure 11. It has to make a decision on which segmented component to connect to in order to have the best chance to return to the same component it starts from along a smooth path without revisiting any segments that have already been traversed. Once the decision is made, it picks one of the endpoints of the next component to enter and exit from the other endpoint.

**Fig. 11.**
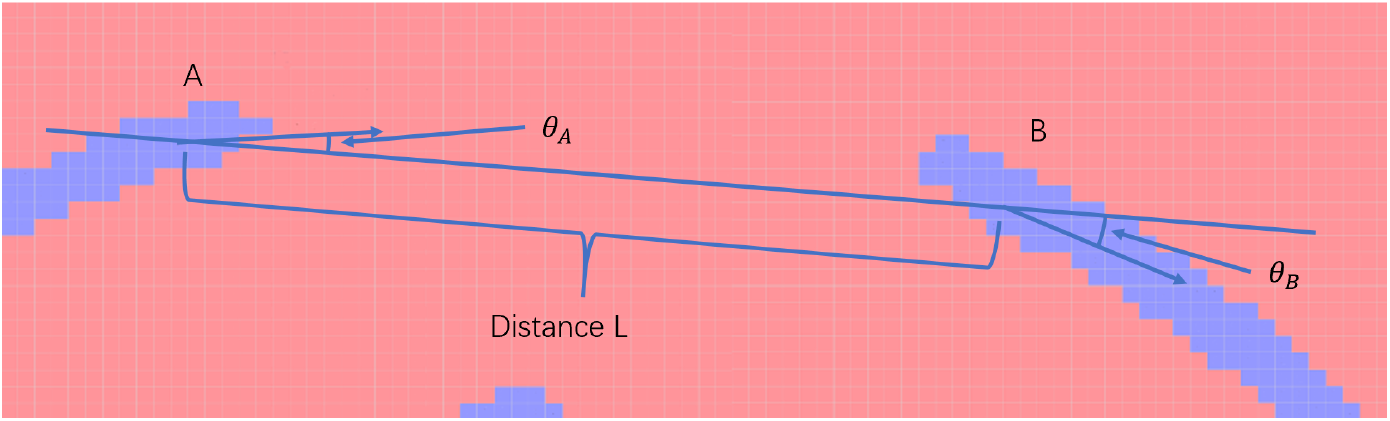
The local reward for moving from *A* to *B* is calculated from the distance L between *A* and *B* as well as the difference between angles *θ_A_* and *θ_B_*.

In this case, the local reward for jumping to an endpoint *B* of another segmented component is determined by the distance between *A* and *B*, the direction of the line segment *AB*, as measured by the angles formed between *AB* and the tangent lines of each component near *A* and *B*, as well as the approximate curvatures of the segments containing *A* and *B* respectively. (See Figure 11)

The *Q*-value of each action is initially determined by the local reward. However, taking the action associated with the highest *Q*-value does not guarantee that the walker can return to the segment it starts from within a fixed number of jumps, which form a *training episode*. Therefore, multiple episodes may be needed to produce a *Q*-table that provides an optimal policy for the walker to successfully return to the original segmented component after going through several components that can be connected in a smooth manner. The process of generating an optimal policy (encoded by a *Q*-table) is often referred to as *Q*-learning. During the training and learning process, entries of the *Q*-table are updated according to the standard Q-learning update formula

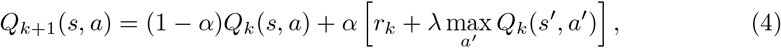

where *k* is the episode number, *r_k_* is the local reward for taking action *a* at *s* in *k*th episode, *s*′ is the observed state that can be reached from s by taking the action *a*, 0 < *α* < 1 is the learning rate and 0 < λ < 1 is the discount rate. For each state, weapply the so-called *ϵ*-greedy policy to select an action. In such a policy, we make the walker take the action that yields maximum *Q*(*s, a*) value most of the time, but allow it to take a random action occasionally. The value of *ϵ* defines the probability of taking a random action in an *ϵ*-greedy policy. To be specific, before taking an action, we generate a random number. If the number is less than *ϵ*, we pick one of the possible actions randomly. Otherwise, we pick the action *a* that yields that largest *Q*(*s,a*) for the current *s*. Both *α* and *ϵ* can be adjusted dynamically. For example, at the beginning of the training episodes, both *ϵ* and *α* are set to a relatively large value. They are gradually decreased in subsequent episodes.

The basic procedure for updating the *Q*-table in a *Q*-learning algorithm is summarized in Algorithm 1 [15]. In this algorithm, *N*_episode_ refers to the number of episodes allowed to identify one set of connected components, max_act_ refers to the maximum number of actions that can be taken within one episode. For simplicity, we assume a fixed learning rate *ϵ* is used. The function `rand()` generates a uniformly distributed random number between 0 and 1.

**Algorithm 1.**
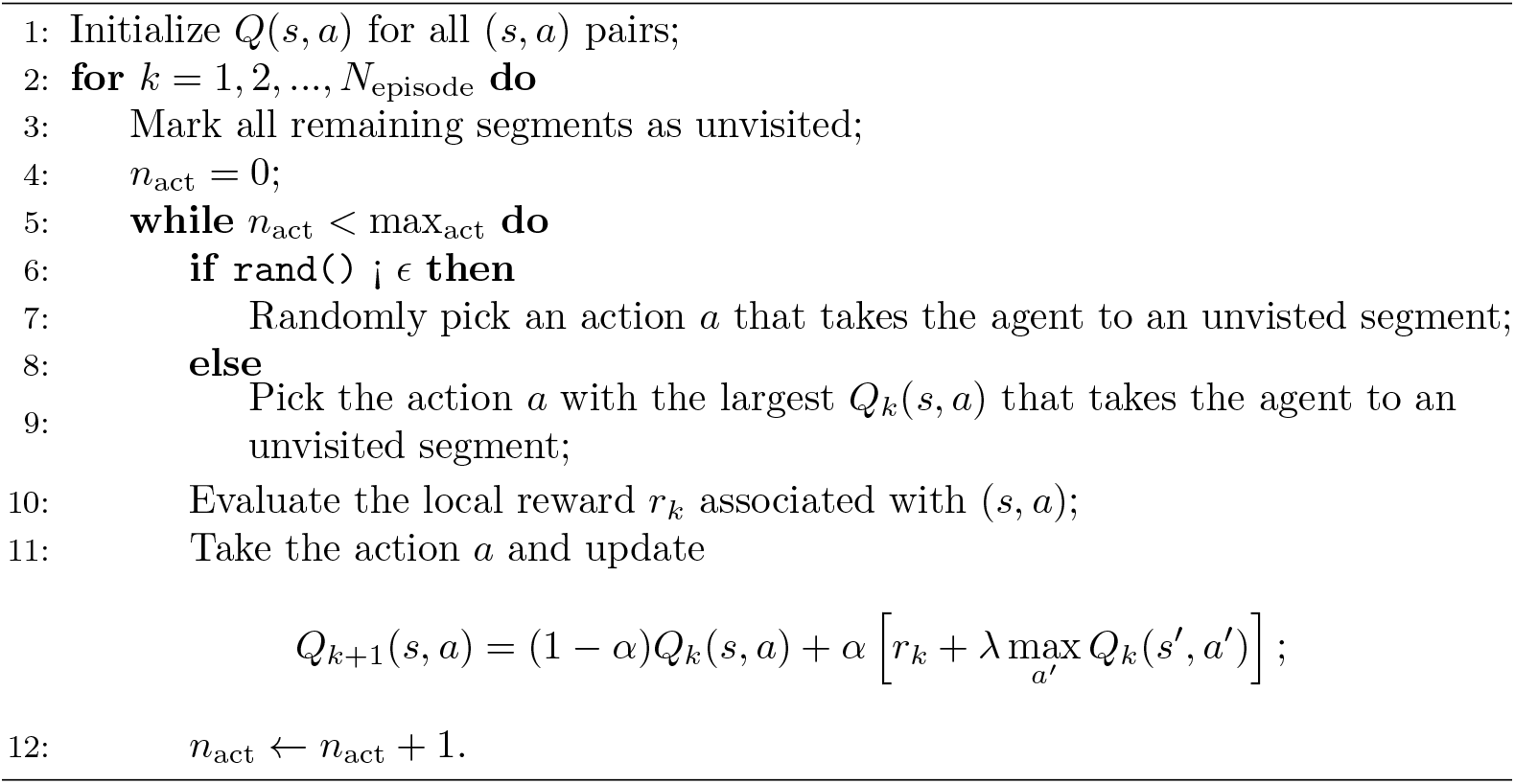
*Q*-learning with *ϵ*-greedy policy

The local reward *r_k_* (in line 10 of the algorithm) we assign to each action of moving from the end of one segment to that of another is typically a negative number determined by the distance of the segments, the difference in their curvature and orientation (in terms of incident angles). The reward is much smaller (more negative) if the transition from one segment to another is not smooth. A positive reward is given only when the action takes the agent (walker) back to the segment it starts from. If no positive *r_k_* is ever generated in an episode, that episode is considered a failed episode.

After a *N*_epsisode_ *Q*-learning step is terminated, we can use the final *Q*-table to define the optimal policy for connecting different components. We start from starting component and connect to the next component by selecting among all actions that yield the largest *Q*(*s,a*) value.

Figure 12(a) shows the final *Q*-table as a heatmap produced from several *Q*-learning episodes applied to the identified segments shown in Figure 10. These episodes start from segment 3 shown in Figure 10. The *Q*-table heatmap is rescaled to make it easier to interpret. The vertical axis labels the states, which are the segment numbers, and the horizontal axis labels the actions the agent can take, which are labeled by the target segment numbers since each action corresponds to a jump from one segment to another. According to this table, if the agent starts from segment 3, the next action will take it to segment 4, followed by another action that takes it back to segment 3. This sequence of actions allows us to connect segments 3 and 4. Figure 13 shows all connections made by the RL algorithm when it is applied to the image shown in Figure 10. Each connection is marked by a black line connecting two segments. Although all connections in this particular example consist of two segments, the RL algorithm we developed can be used to connect several segments. Figure 14(a) shows a heatmap representation of the *Q*-table produced by the RL algorithm when it is applied to multiple hand-drawn segments shown in Figure 14(b). Using the *Q*-table, we can easily connect segments labeled by 3,7,2,6,0,8,1.

**Fig. 12.**
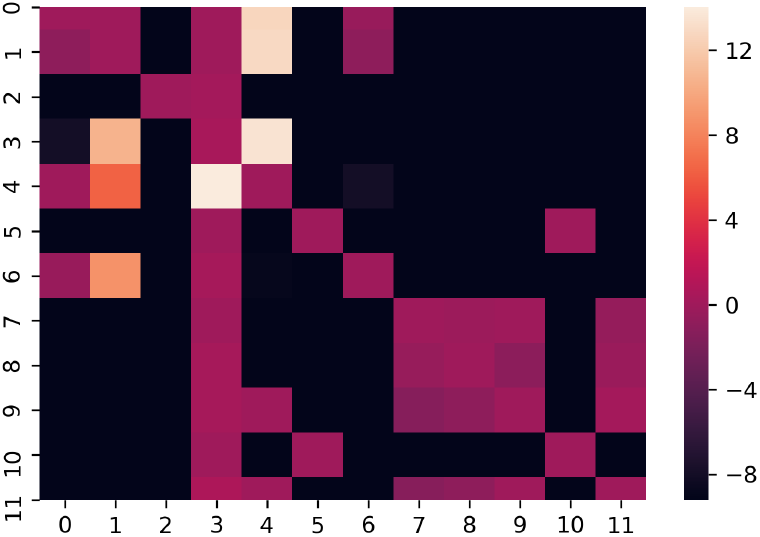
The heatmap produced from several episodes of the RL algorithm applied to Figure 10 starting from segment 3

**Fig. 13.**
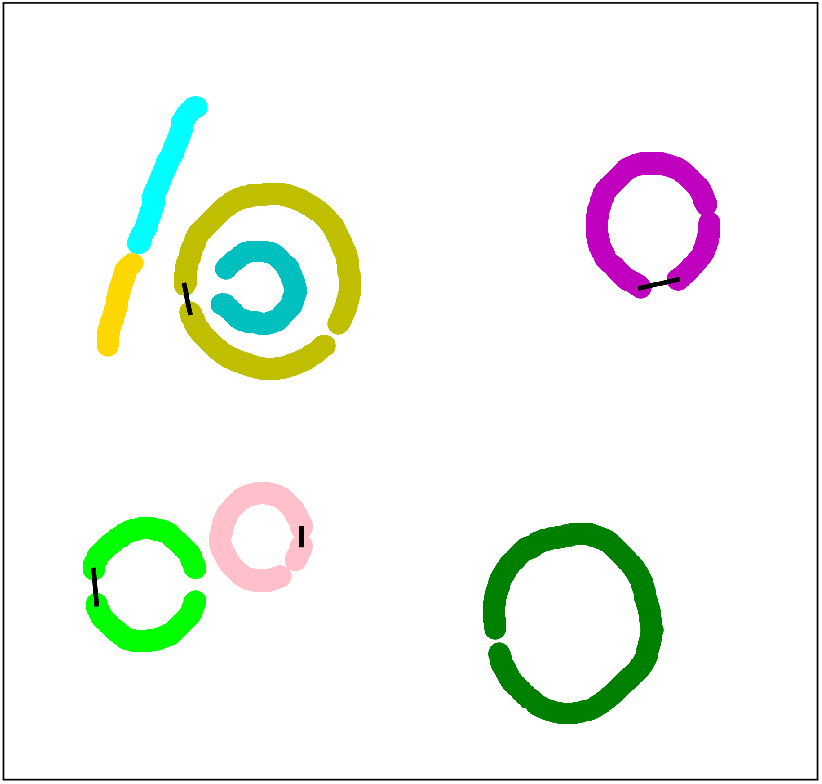
All connections made by the RL algorithm. Each connection is marked by a black line in the figure.

**Fig. 14.**
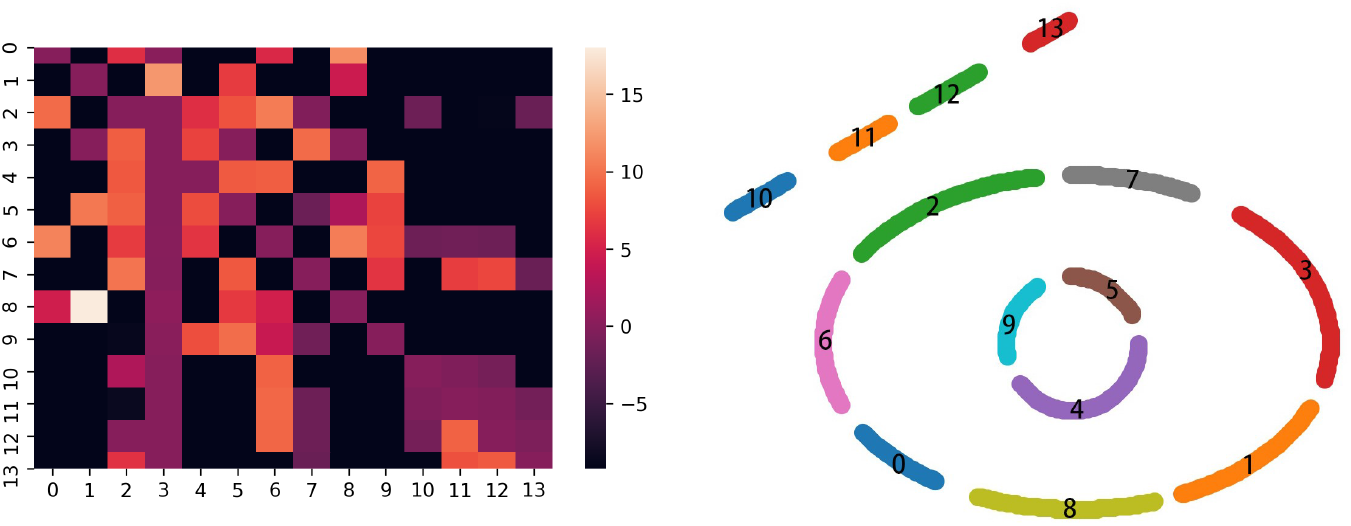
(a) The heatmap representation of the *Q*-table produced from several episodes of the RL algorithm applied to connect the hand drawn segments in (b). The connected path is [3,7,2,6,0,8,1].

### 5.2 Classification via Region Merging

What is essentially achieved by the reinforcement learning algorithm presented above is the classification of U-Net segmented components into separate classes that represent membrane structures of distinct subcellular structures. Segmented components belonging to the same class can be connected via a number of geometric fitting procedure to be discussed in section 5.3.

In this section, we consider another strategy to perform this classification. Standard classification techniques such as K-means and principal component analysis are not directly applicable because the objects to be classified in our case are ordered sets of pixels with different sizes. They are not convenient descriptors as are used in the standard classification methods which treat each data point as a vector of numbers.

The classification scheme we use is a variant of the statistical region merging method originally developed in [16]. In this approach, each pixel identified by U-Net to be part of the membrane forms its own region initially. Regions that are sufficiently close are merged successively. The distance between two regions *R*_1_ and *R*_2_ is defined by

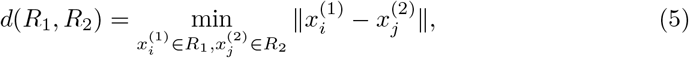

where 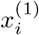 and 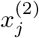 are the coordinates of two points in regions *R*_1_ and *R*_2_ respectively. In addition to using *d* as a metric to decide whether to merge two adjacent regions, other visual cues such as curvature of the existing region can be used to define a predicate for reaching a merging decision. Such a predicate may also take into account uncertainty in the data (due to the presence of noise, artifacts and missing information) to allow merging decisions to be made on a statistical basis [17].

At the end of the merging process, each distinct region represents a distinct (membrane) class which is assigned a unique label.

If we simply perform region merging within each 2D tomogram slice, disconnected components with a relatively large gap will remain in different regions and thus be disconnected after the merging process is completed. However, if we allow merging to be performed in 3D, i.e., allowing pixels in different slices to be merged into the same region, then two disconnected segments on the same membrane can be merged into the same region when each one of these segments contains separated pixels that can be connected to other pixels in an adjacent slice that have already been merged into a common region. This is possible because segmented components that belong to the same membrance surface may be disconnected at different locations in different slices. By exploiting the continuity of a membrane structure among different tomogram slices, we can successfully place two disconnected segments in a single tomogram slice into the same class.

Figure 15 shows that even though segments A and B are disconnected in slice 90, they contain pixels that can be connected (via the path shown in the right subfigure) to other pixels in an adjacent slice that has been merged into a common region on slice 105.

**Fig. 15.**
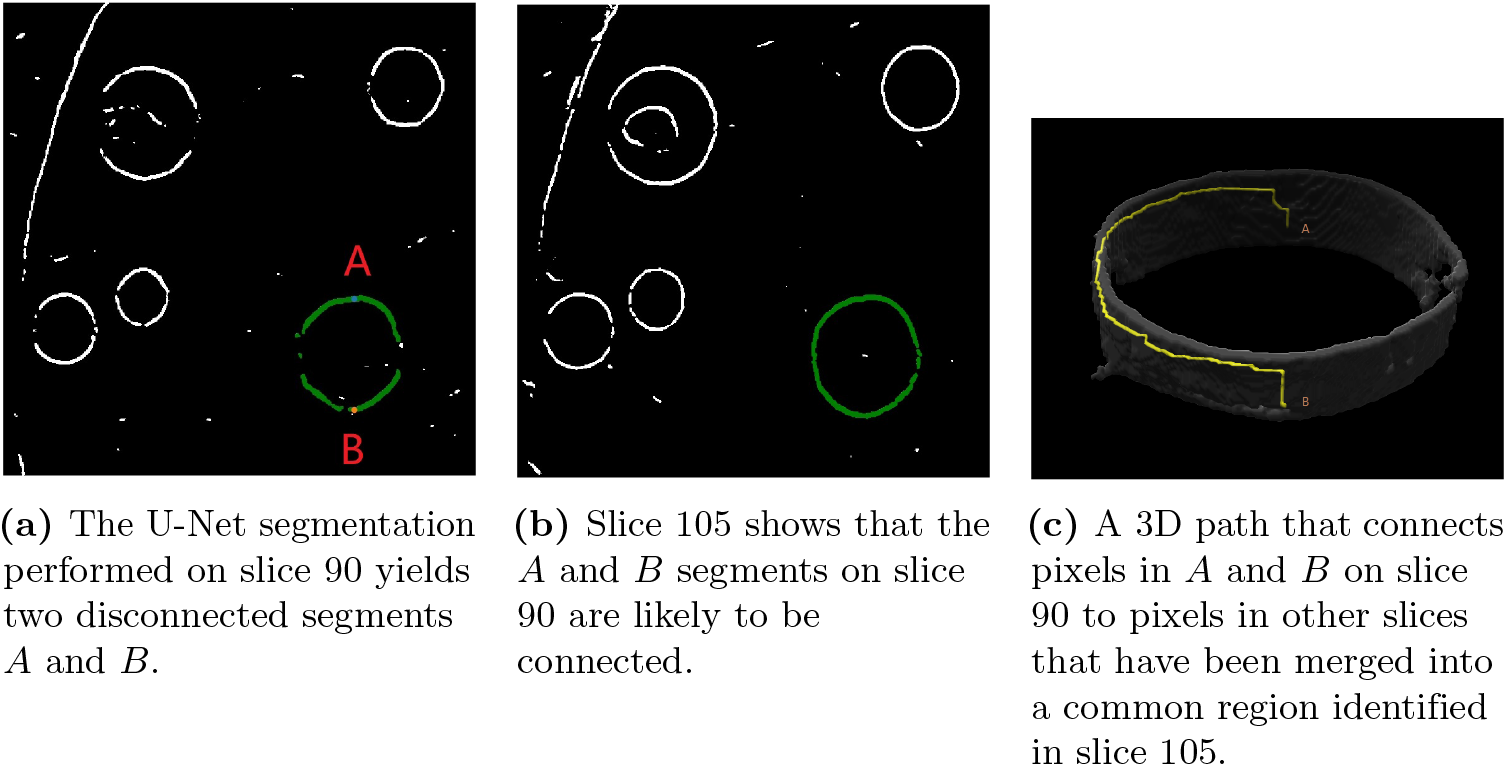
Placing two disconnected segments *A* and *B* in slice 90 into the same class through 3D region merging.

To accelerate the class merging process, we use an efficient union-find data structure and a tree-based merging algorithm [18]. In such an algorithm, each pixel is treated as a node on a tree. Pixels merged into the same class are nodes on the same tree organized in a hierarchical fashion. Each tree is labeled by its root node. Finding all pixels belonging to the same tree essentially amounts to a breadth-first traversal of the tree, which is constructed as it is being traversed. We can start from an arbitrary pixel and add all adjacent pixels as its children in a tree. Each of its children adds more descendants to the tree until no pixel can be added. If there are still pixels that have not been used, the construction of a new tree starts from an arbitrary unused pixel. This process continues until all pixels are used.

An alternative way to construct these trees simultaneously is to go through all pixels in some order. A pixel *B* adjacent to the pixel *A* being examined is added as a child of *A* in the tree *T_A_* that *A* belongs to if *B* has not been merged with other pixels in another tree. Otherwise, the tree that contains *B*, denoted by *T_B_*, is merged with *T_A_*. The merge involves placing the root of *T_B_* as a child of *A* in *T_A_*. Once we have gone through all pixels, each pixel will belong to one of the trees. To find out which tree each pixel belongs to, we simply traverse from the node the pixel is mapped to towards the root since the root node is essentially the label of the tree.

Figure 16 shows the final six regions created by the region merging procedure. We assign a different color for each region, which corresponds to one membrane structure (except the sheet in the upper left corner of the 3D rendering.) Although these regions can be viewed as a 3D segmentation of the tomogram, the segmented structures contain visible artifacts such as extra voxels protruding from a membrane surface and gaps in the membrane surface. We will show in section 6 that this problem can be fixed by a 3D fitting and refinement procedure.

**Fig. 16.**
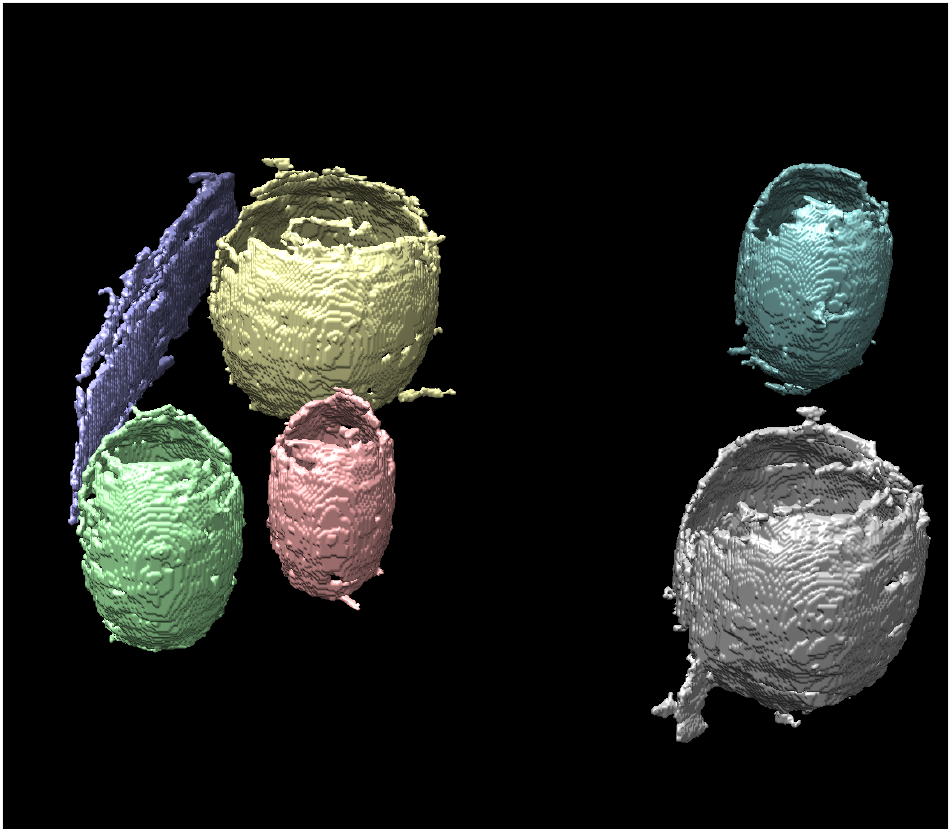
3D rendering of six regions obtained at the end of the region merging procedure applied to the entire tomogram. Each region is labeled by a unique color.

**Fig. 17.**
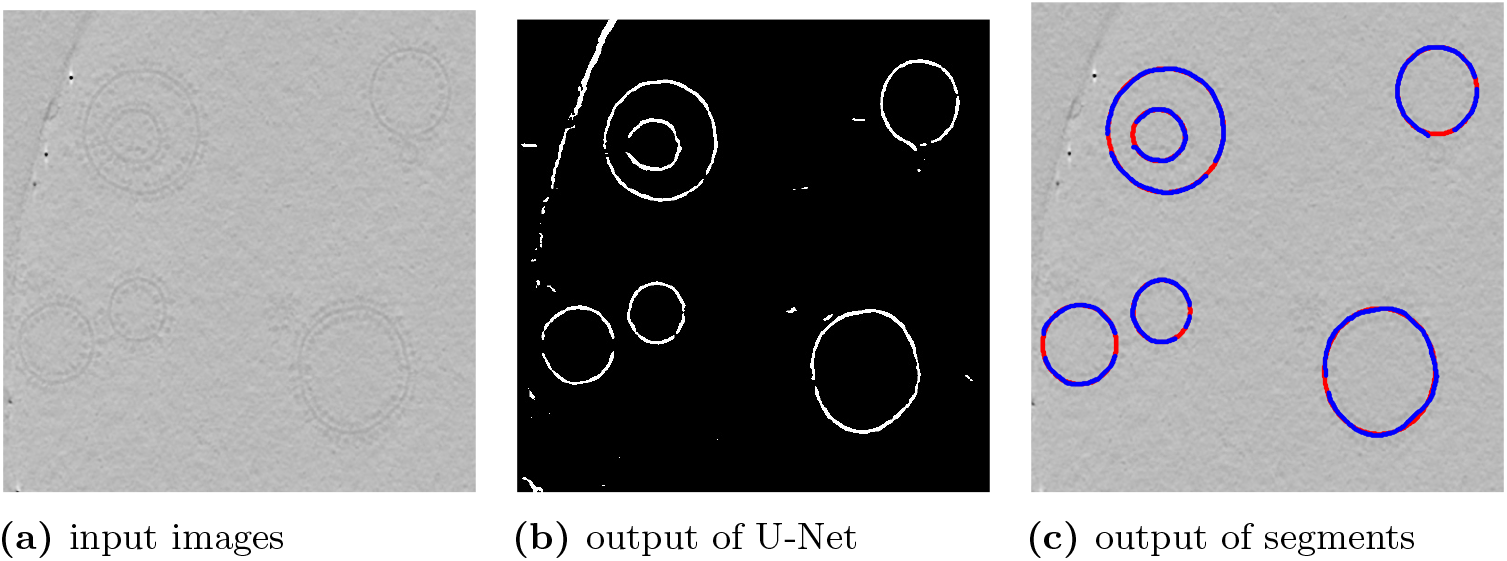
Fitting U-Net segmented components with ellipses for the 120th tomogram slice.

### 5.3 Connecting segmented components via 2D parametric and non-parametric fitting

Once the segmented components have been classified, the pixels belonging to the same class can be connected in 2D via a parametric and non-parametric fitting scheme by taking into account prior knowledge of the subcellular structure to be examined.

#### 5.3.1 Parametric fitting

If the object to be segmented has a simple geometry, we can use a parametric fitting scheme to deduce the missing pieces between disconnected components that have already been segmented out by U-Net. If, for example, the horizontal slice of vesicle membranes in Figure 8(a) all have a elliptical shape, we can parametrize the points on the vesicle membrane as solutions of

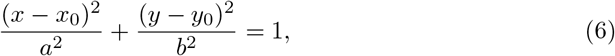

where *x*_0_, *y*_0_, *a* and *b* are parameters to be determined from a nonlinear least squares fitting procedure that minimizes the discrepancy between the left-hand and the right-hand sides of (6) among all pixels on the segmented components that belong to the same class.

#### 5.3.2 Non-parametric fitting via Gaussian process

When the membrane of the subcellar structures cannot be easily described by simple geometric objects that admit an analytic parameterization, we use a non-parametric fitting procedure based on the Gaussian process (GP) formalism [19] and the implicit surface [20] formulation.

The basic idea is to view the 2D curve that encloses a subcellular structure as the zero level set of a smooth 2D scalar function *f*(*x,y*). Our goal is to construct this non-parametric function *f*(*x,y*) such that *f*(*x,y*) = 0 for (*x,y*) ∈ *U*, where *U* contains pixels in segmented components that have been identified and connected by algorithms presented in sections 4, 5.1 and 5.2. In addition to the segmented pixels, the set *U* also includes the pixel coordinates of a number of anchor points both inside and outside of the expected membrane surface so that a smooth convex or concave function *f*(*x,y*) can be constructed. The choice of these anchor points represents our prior belief that certain parts of the image should belong to the exterior of the membrane while the other parts should belong to the interior even though we do not know the precise location of the interior/exterior separation in the region of interest.

We set the values of *f* to negative and positive constants at these anchor points as shown in the example given in Fig. 18.

**Fig. 18.**
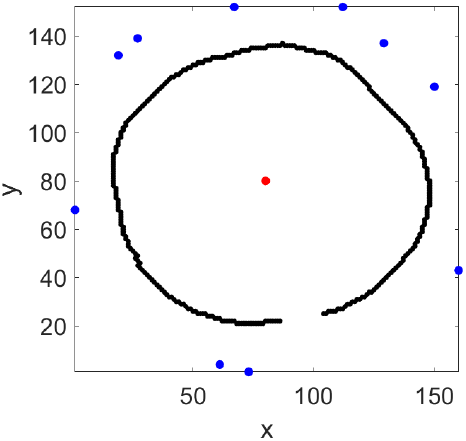
The anchor points added to a partially segmented membrane slice. The value of *f*(*x,y*) is set to −1 for blue anchor points (outside the membrane), and to 1 for the red anchor points (inside the membrane).

If *f*(*x, y*) is continuous and sufficiently smooth, the pixels in the zero level set of *f*(*x, y*) that have not been included in the set *U* defined above will fill in the gaps of the partially segmented membrane components returned from algorithms used in sections 5.1 and 5.2 to form a continuous and smooth boundary (surface) as shown by the example given in Fig. 19.

**Fig. 19.**
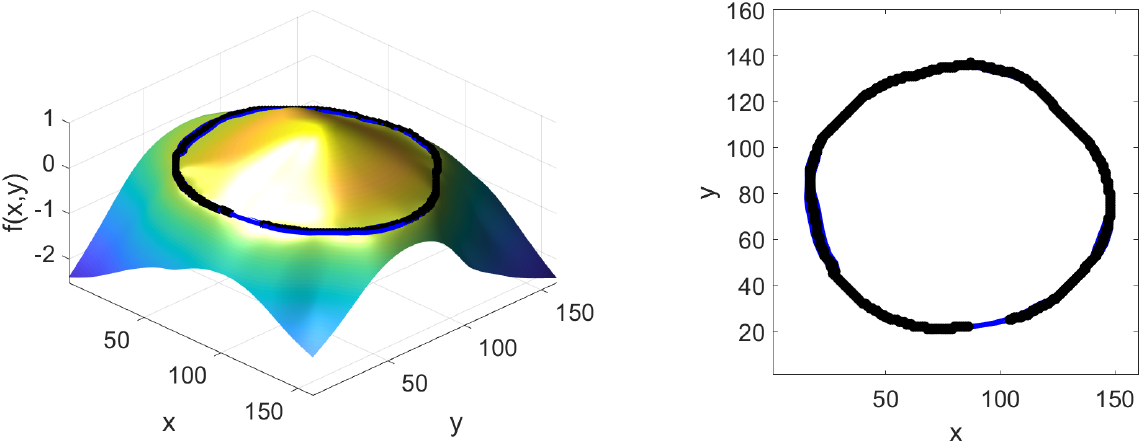
The mean function produced by GP (left) and its zero-level set (right).

A GP is a prior of the distribution of functions *f* that is generally defined by a multivariate Gaussian 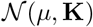, with a mean function *μ* and covariance function **K**. Since we are only interested in function values at the *n* pixels of a 2D image, *μ* is a vector of length *n*, and **K** is a *n* × *n* matrix. We denote the function values of *f* on *n* pixels by **f**.

The vectors **f** and *μ*, and the covariance matrix **K** can be partitioned as

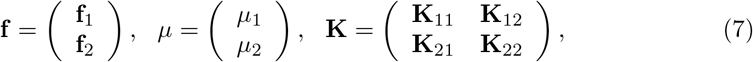

where **f**_1_ corresponds to random variables associated with values of *f* defined on pixels contained in the set *U* described above, which includes both the coordinates of the segmented components and the coordinates of the anchor points, and **f**_2_ corresponds to random variables associated with values of *f* defined on the other pixels in the image. The vectors *μ_i_* are the means of **f**_*i*_, *i* = 1, 2 respectively. The partition of **K** is conformal to that of **f** and *μ*.

The conditional probability density function (PDF) of **f**_2_ given **f**_1_ yields the posterior PDF of **f**_2_ given **f**_1_. It is well known [21] that this PDF is a multivariate Gaussian also with the mean

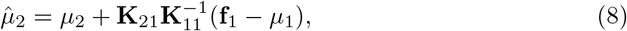

and covariance matrix

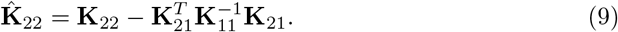

The mean 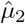 yields a good estimate of the values of *f* on pixels outside of *U*. It allows us to reconstruct the missing components on the membrane surface by finding pixels (*x,y*) ∉ *U* that satisfy 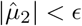 for some small constant *ϵ*. In practice, *μ*_1_ and *μ*_2_ are often set to 0. Therefore, (8) and (9) can be computed explicitly through the solution of a linear system, matrix-vector and matrix-matrix multiplications. Regularization may be needed when **K**_11_ is ill-conditioned. Fig. 19 (left) shows the mean function defined on pixels outside of the segmented surface (curve) shown in Fig. 18. The segmented surface is shown in black. The zero level set that fills in the opening on the segmented surface is shown in blue. The figure on the right shows more clearly the reconstructed surface (curve) as a 2D contour.

In addition to providing a mean estimate of where the missing components of the segmented surface (curve) should lie, we can also quantify the uncertainty associated with the reconstructed surface by evaluating the marginal likelihood of a pixel being on the zero level set of *f*, i.e.

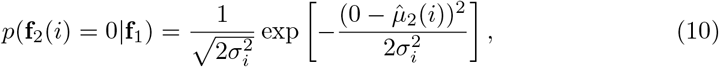

where **f**_2_(*i*) and 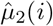 denote the *i*th component of **f**_2_ and 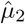 respectively, and *σ_i_* is the ith diagonal element of 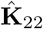. This marginal PDF quantifies the uncertainty of a particular pixel being on the surface of the membrane. Fig. 20 shows the marginal PDF associated with the 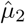 shown in Fig. 19 as a grayscale image. The darker the pixel, the higher the likelihood of the value of 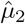 being zero (hence on the membrane) at that pixel. We exclude the previously segmented pixels by setting the color of these pixels to red.

**Fig. 20.**
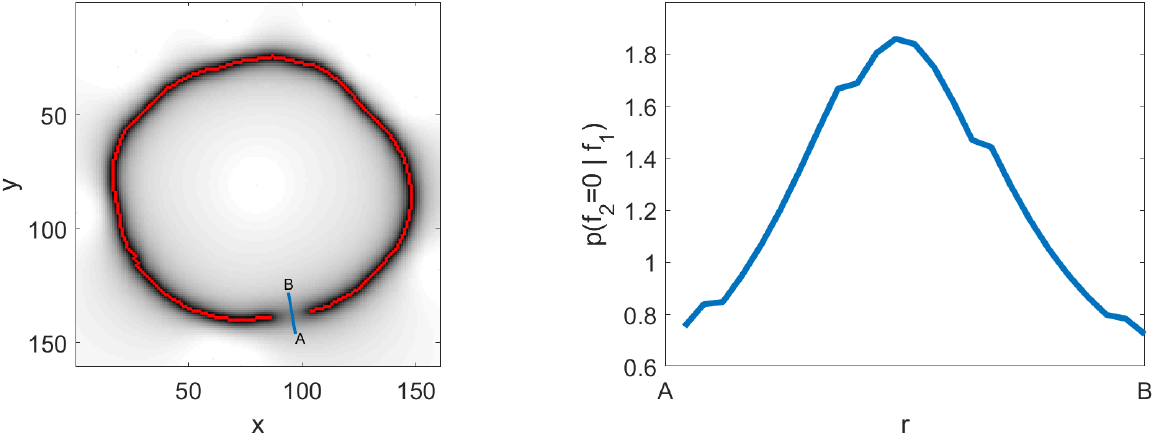
The marginal likelihood of each pixel being on the zero level set of *f*, i.e. on the surface/boundary of the membrane (left). The marginal likelihood of pixels along the line segment AB (shown in the left figure) being on the zero level set of *f* (right).

Note that the GP prior on the distribution of **f** is largely determined by the covariance **K**. The mean *μ* does not play an essential role and is usually set to 0. The covariance describes how function values *f*(*x,y*) are correlated for different (*x,y*)’s. It is often expressed in terms of the distance between different (*x,y*)’s. A commonly used covariance kernel is the Gaussian kernel defined as

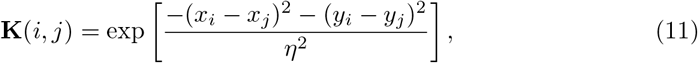

for some appropriately chosen length scale parameter *η*, where (*x_i_, y_i_*) and (*x_j_, y_j_*) are coordinates of the *i*th and *j*th pixels respectively.

However, this particular kernel does not work well in sufficiently constraining the zero level set of *μ*_2_ by that of *μ*_1_ through the smoothness of *f*. A more effective covariance kernel proposed in [22] has the form

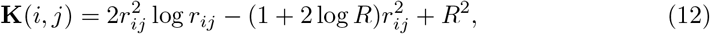

where 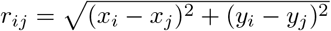 and *R* is the maximum distance between any two pixels in the 2D image. Note that *K*(*i,i*) = *R*^2^ for all *i*. This kernel function is the Green’s function of a 4th order differential operator. It is related to smoothing splines interpolation [23] and the thin-plate spline regularizer [22].

The GP framework is flexible in allowing us to construct the mean of *f* (and the implicit surface associated with its zero level set) to match prior knowledge about certain biological structures. For example, by placing one anchor point within the inner ring of the double membrane structure present in, for example, Fig. 17(b), some anchor points between the inner and outer rings and some outside of the outer ring, and setting the values of anchor points to −1, 1 and −1, we can reconstruct the missing segments in both the inner and outer membrane as can be seen in Fig. 21.

**Fig. 21.**
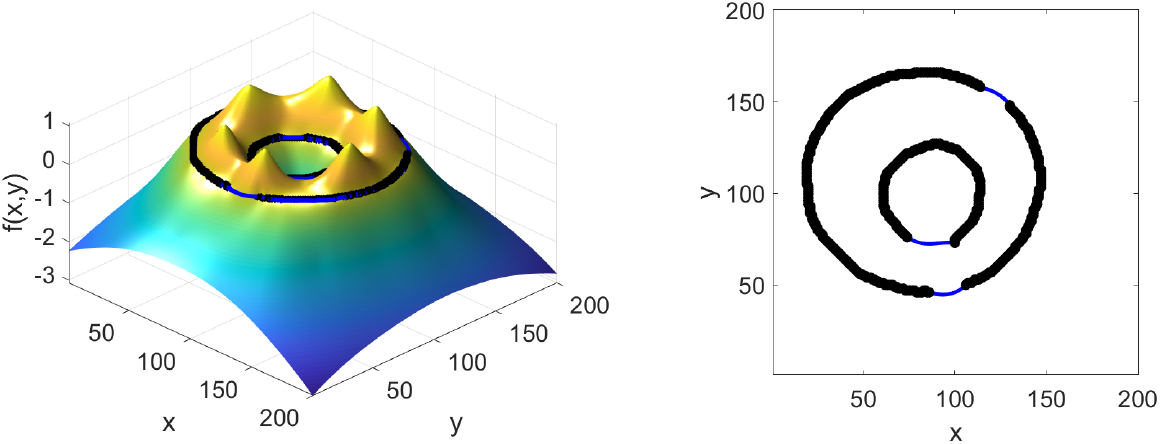
The mean function produced by GP and the zero-level set for inner and outer surfaces.

## 6 Refinement in 3D

Although 2D parametric and non-parametric fittings allow us to fill in the missing pixels in the segmented component in each slice of the tomogram and produce smooth 2D curves in each slice, stacking these 2D curves together may produce a nonsmooth 3D surface with gaps or bumps along the vertical direction as can be seen in Figure 22.

**Fig. 22.**
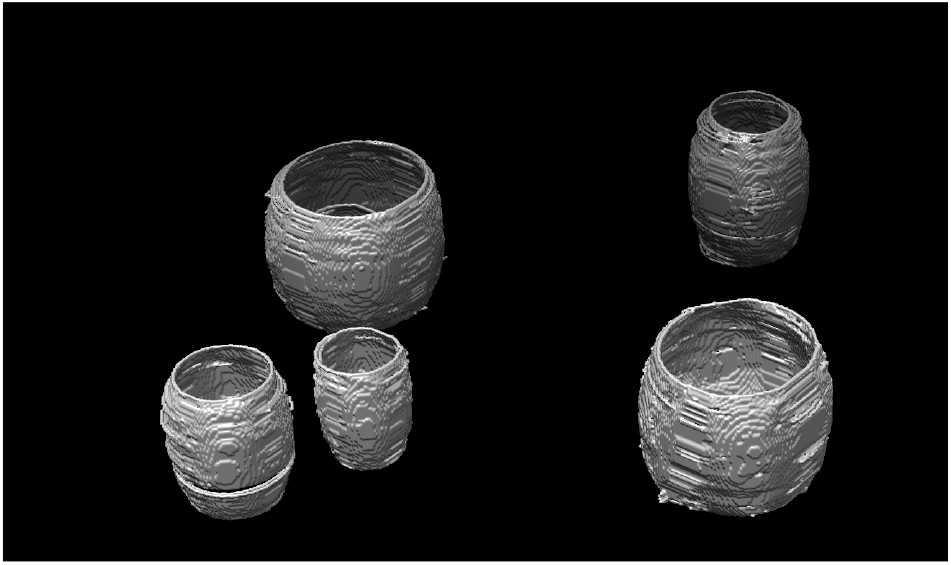
The isosurface of the 3D segmentation obtained by stacking the segmented slices 60 through 160.

The nonsmoothness of the surfaces can also be quantified by the maximum of distances between each membrane pixel and its nearest neighboring pixel on an adjacent slice. We plot in Figure 23 a histogram of such maxima for all slices shown in Figure 22. We can see from the histogram that most of the maximum distances are within 2 pixels, but there are quite a few between 2 and 5 pixels. There is even one that is 8 pixels long.

**Fig. 23.**
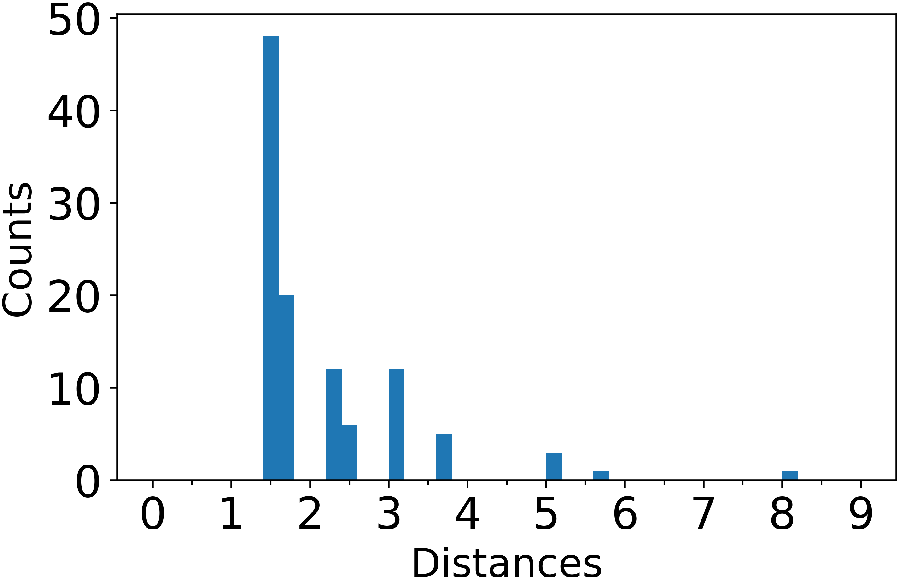
Histogram of the maximum distance between a membrane pixel and its nearest neighboring membrane pixel on an adjacent slice for all slices shown in Figure 22.

The lack of smoothness along the vertical direction results partially from artifacts produced by the U-Net segmentation which picks up some spurious pixels that do not belong to the surface of the membrane. It may also be caused by either an ill-posed 2D fitting (due to the presence of only a few pixels grouped into the same class) or overfitting that tries to connect pixels on the membrane with spurious pixels. Neither procedure is properly constrained by the continuity and smoothness of the membrane surface across tomogram slices.

To address this problem, we develop a refinement procedure to first collect segmented pixels in different tomogram slices that belong to the same membrane surface. Spurious pixels are pruned. We then use the 3D Gaussian process formalism to construct 3D membrane surfaces that are zero level sets of a continuous function defined on a 3D volume and anchored by a few voxels both inside and outside of the membrane surfaces to be reconstructed.

### 6.1 Voxel selection

In order to make effective use of the Gaussian process technique in 3D to construct the desired membrane surfaces, we need to identify as many voxels that lie on the same surface as possible. These voxels are collected from segmented pixels within each tomogram slice. Pixels that belong to the same class produced from the classification schemes discussed in sections 5.1 and 5.2 are grouped together. However, in some tomogram slices, only a few pixels belonging to the surface are visible. Even fewer can be identified by the U-Net. Although parametric or non-parametric fitting can be used to reconstruct some of the pixels, the lack of visible pixels in these slices makes the fitting procedure ill-posed. For example, Figure 24 shows that U-Net picked up a few pixels on the inner membrane of a vesicle in the upper left corner of the 100th tomogram slice. Some of these pixels were filtered out during the classification procedure because they are isolated and not connected to other pixels. Only a small number of pixels at the top of the inner membrane are retrained. When an elliptic parametric fitting procedure is applied, a small ellipse is produced, which does not correctly characterize the shape of the inner membrane.

**Fig. 24.**
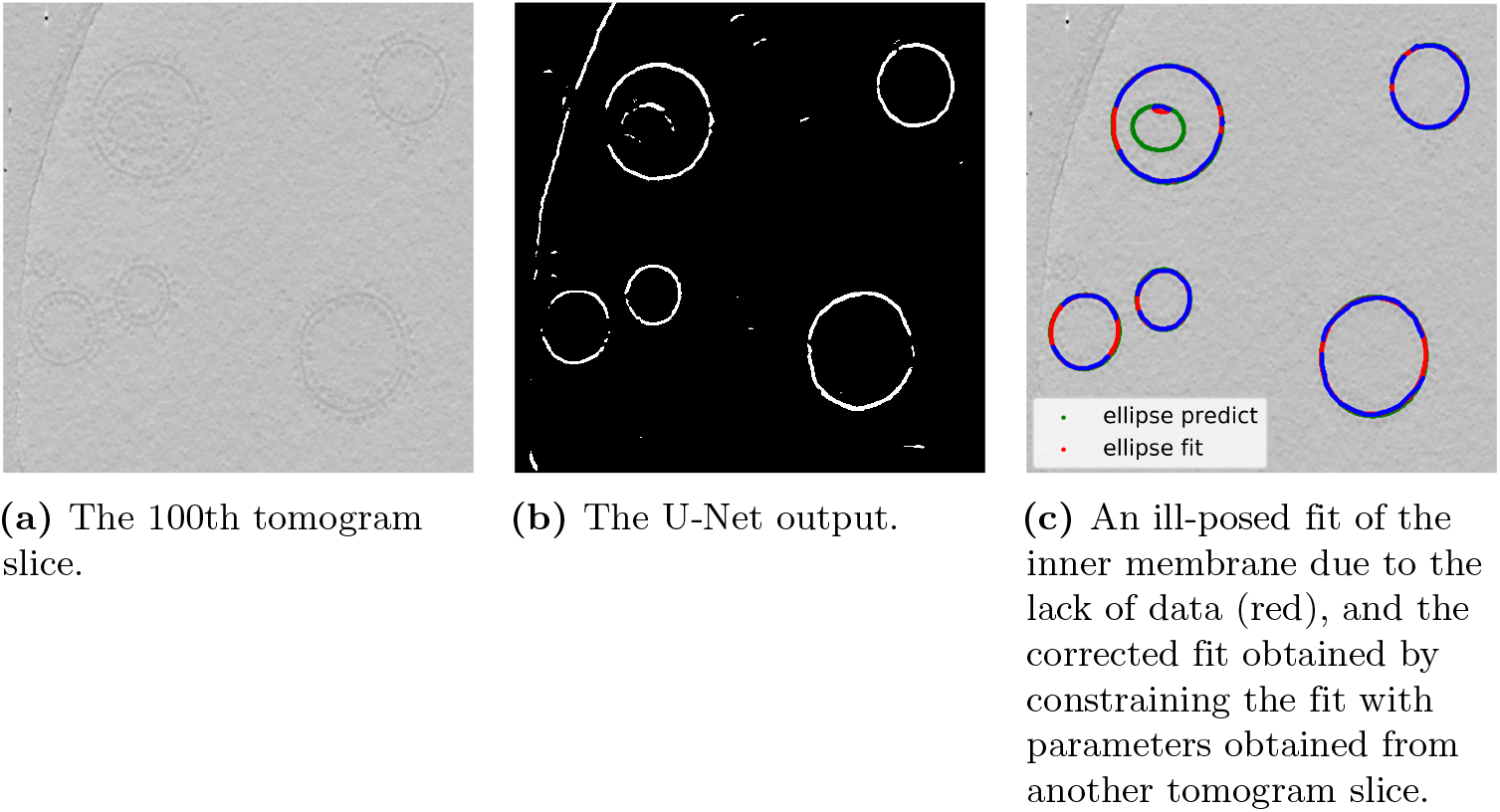
An elliptic fitting of the inner membrane on the 100th tomogram slice using filtered U-Net segmentation and the correction made by taking into account the fitting made on the 120th slice.

However, the correct shape of the inner membrane is obtained when the parametric fitting procedure is applied to the 120th tomogram slice shown in Figure 17. For that tomogram slice, many pixels can be seen to lie on the inner membrane. They are correctly identified by the U-Net segmentation. The parameters associated with the ellipse that closes the gap in the U-Net segmented inner membrane can be used to train the parameters to be optimized when a nonlinear least squares fitting is applied to the 100th tomogram slice. To be specific, we can use the parameters *x*_0_, *y*_0_, *a* and *b* obtained from the least squares fit of the inner membrane for the 120th tomogram slice as the starting guesses to the parameters associated with the ellipse that fit the selected pixels in tomogram slice 100. Upper and lower bounds for these parameters are also set based on the parameters obtained from the 120th tomogram slice. The constrained optimization with a good starting guess yields an ellipse shown as the green curve in Figure 24(c). Once this ellipse is constructed, all segmented pixels produced by U-Net that are sufficiently close to the ellipse are selected as voxels to be used in the 3D Gaussian process fit.

### 6.2 3D fitting via Gaussian process

Once all valid voxels for each one of the organelles in the tomogram have been identified, we use the technique of Gaussian process discussed in section 5.3.2 to construct an isosurface that connects all these voxels. This isosurface is defined as the zero level set of a 3D function *f*(*x, y, z*) that is smooth. For an organelle with a single membrane, we choose an anchor point interior to membrane and set the function value of that point to 0. This point can typically be chosen as the centroid of all validated voxels that are considered to be on the membrane. A number of anchor points (*x_i_, y_i_, z_i_*) in the exterior region of the membrane must also be chosen. The value of *f* is set to 1 at these anchor points. There are a few ways to choose these anchor points. These choices represent our prior knowledge of the shape of the membrane. For example, if the organelle is believed to have an ellipsoidal shape, we can enclose the validated voxels associated with this organelle by an ellipsoid with an appropriate size and orientation estimated from the selected voxels, and sample quasi-uniformly on the surface of the ellipsoid. Another possible way is to simply choose a few validated voxels that are well separated, and extend the ray connecting the centroid with the selected voxel proportionally to the distance between the selected voxel and the centroid (See Figure 25). For example, point *A* is obtained by connecting the centroid at *C* with a validated voxel *B* and extending the ray so that the distance between *A* and *C* is 1.2 times the distance between *B* and *C*.

**Fig. 25.**
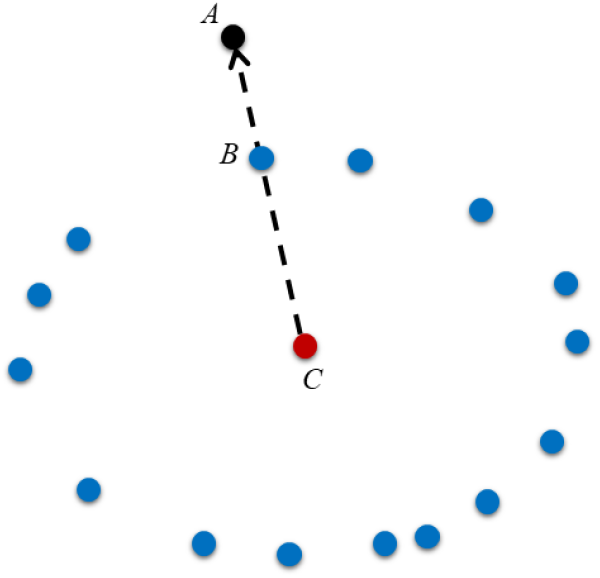
The anchor point (*C*) is chosen by connection a validated voxel *B* with the centroid of all validated voxels *C* and extending the ray away from the centroid so that |*AC*| = 1.2 × |*BC*|.

For 3D fitting, each element of the covariance matrix associated with the joint Gaussian distribution (11) is chosen as

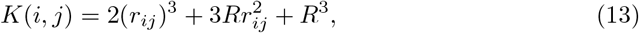

where 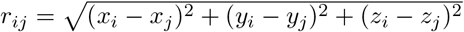, and *R* is the maximum distance between any two validated voxels that are considered to be on the membrane surface of a single organelle.

Figure 26 shows the lower half of the reconstructed surfaces for several organelles within the tomogram as well as the validated voxels selected for fitting in the upper half of the tomogram.

Figure 27 shows the entire reconstructed exterior surfaces that characterize the shape of these organelles as well as the ATP synthase proteins identified by the U-Net.

**Fig. 26.**
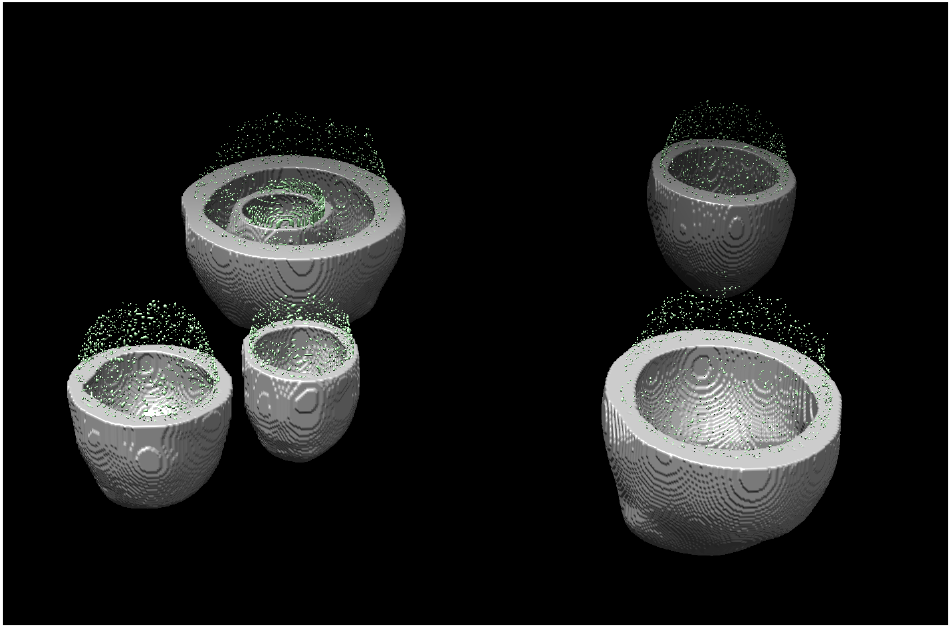
Lower half of the reconstructed membrane surfaces for several organelles within a single tomogram as well as validated voxels selected for the GP fitting procedure.

**Fig. 27.**
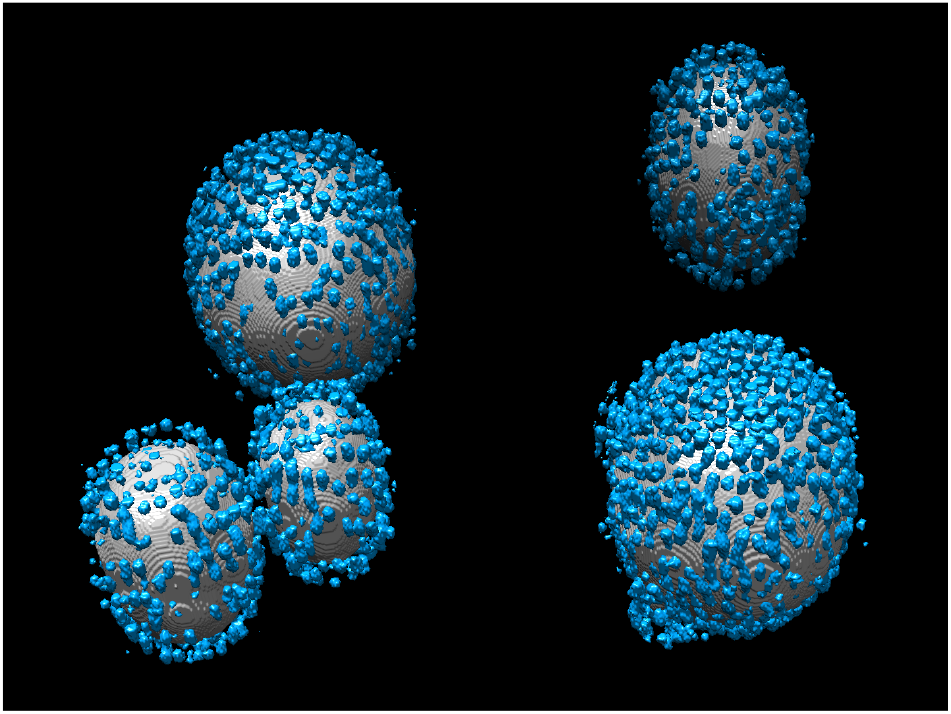
The entire exterior surfaces of the reconstructed membranes of different organelles within the tomogram as well as the ATP synthase proteins (blue) outside of the membranes.

## 7 Discussion and Conclusion

The extreme low contrast of cryo-electron tomograms and artifacts introduced by the limited sample tilt range that is accessible during imaging (the missing wedge problem) makes it difficult to use existing segmentation tools developed in the last few decades mainly for high contrast 3D medical imaging to analyze the tomogram and identify important biological structures.

We presented a machine learning-based segmentation approach to overcome this difficulty. Our approach uses a variety of techniques organized in a learning pipeline to automate the segmentation process. The learning pipeline starts from supervised learning via a U-Net trained with simulated data. It continues with semi-supervised reinforcement learning and/or a region merging techniques that try to piece together disconnected components that should belong to the same subcellular structure. A parametric or non-parametric fitting procedure is then used to enhance the segmentation results and quantify uncertainties in the fitting. Domain knowledge is used in generating the training data for U-Net and in guiding the fitting procedure through the use of appropriately chosen priors and constraints (e.g., anchor points for GP). We demonstrated that the approach proposed here worked well for extracting membrane surfaces of protein reconstituted liposomes in a cellular environment that contains other artifacts. Although we have only demonstrated the effectiveness of our approach on one dataset, the approach itself is quite flexible and can be applied to a different dataset with minimal modifications. New domain knowledge for a different dataset can be incorporated by providing new simulated training data for U-Net and new priors through the choice of different anchor points in GP fitting.

## Acknowledgement

This research was supported by the Laboratory Directed Research and Development (LDRD) Program award 20-122 of Lawrence Berkeley National Laboratory (LBNL) under U.S. Department of Energy Contract No. DE-AC02-05CH11231. Z. Li would like to acknowledge support from both LBNL and the Chinese Ministry of Education for carrying out this research during his visit to LBNL.

